# Reducing CETP activity prevents memory decline in an Alzheimer’s disease mouse model

**DOI:** 10.1101/2024.11.05.621935

**Authors:** Jasmine Phénix, Isabel Sarty, Megan S. Katz, Hannah Nie, Anja Kerksiek, Robert S. Kiss, Dieter Lütjohann, William A. Pastor, Judes Poirier, Lisa Marie Munter, PREVENT-AD Research Group

## Abstract

Epidemiological studies have shown that lower activity of the cholesteryl ester transfer protein (CETP) correlates with reduced Alzheimer’s disease (AD) risk. While small molecule CETP inhibitors like evacetrapib have previously been assessed for cardiovascular diseases, their involvement in AD has not been investigated. Here, we establish CETP as a novel pharmacological target for AD treatment. Using CETP transgenic mice crossed to a mouse model of amyloidosis and administering evacetrapib, we provide evidence that CETP inhibition maintained memory independent of classic AD markers, increased hippocampal cholesterol, altered plasma lipoproteins, and changed transcription of genes linked to brain barriers. Using proteomic data of cerebrospinal fluid from cognitively unimpaired individuals at risk for AD (the PREVENT-AD cohort), we confirm that our mouse model reflects physiological changes in pre-symptomatic human subjects. We propose the repurposing of CETP inhibitors as an effective therapeutic strategy to delay or prevent cognitive impairment in AD.

## Introduction

The cholesteryl ester transfer protein (CETP) is a plasma glycoprotein primarily secreted by the liver but also present in the central nervous system (CNS) in humans*^1–3^*. The enzyme facilitates the bidirectional exchange of neutral lipids - cholesteryl esters and triglycerides - between lipoproteins in plasma, including pro-atherogenic low-density lipoproteins (LDL) and anti-atherogenic high-density lipoproteins (HDL), ultimately increasing LDL-cholesterol levels*^4, 5^*. Consequently, the protective effects of reduced CETP activity against atherosclerosis and cardiovascular events have been extensively investigated over the past decades*^6–9^*. This line of research led to the development of small molecule CETP inhibitors*^10–14^*. Despite the safety and effectiveness of novel CETP inhibitors in raising HDL-cholesterol, clinical trials failed to demonstrate superior cardiovascular benefits when compared to statins, causing a halt in research on this class of compounds*^10, 11, 13, 14^*. However, new evidence has highlighted the impact of CETP activity on several other diseases distinct from cardiovascular diseases such as type 2 diabetes mellitus, cancer, and neurodegenerative diseases including Parkinson’s disease, Lewy body dementia, and Alzheimer’s disease (AD)*^15–17^*. Additionally, we and others have shown that CETP inhibitors enter the brain, demonstrating versatility in targeting both peripheral and brain tissue*^18, 19^*. As such, these findings have reignited interest in the therapeutic potential of CETP inhibitors.

Several lines of evidence support the hypothesis that reducing CETP activity will preserve memory in AD. Firstly, epidemiological studies comparing centenarians to the general population have revealed that CETP variants with reduced activity are linked to longevity, preserved cognitive function and a lower risk of dementia*^20–22^*. Intriguingly, low CETP activity overrides detrimental effects of the apolipoprotein E4 allele (*APOE4*)*^23–27^*, the greatest genetic risk factor for AD*^28, 29^*. Given that 40-65% of all AD patients are *APOE4* carriers, CETP inhibition holds great potential as a novel therapeutic target in AD research*^30^*. Secondly, CETP has a specific function as a non-promiscuous enzyme and thus constitutes an excellent candidate for pharmacological inhibition*^1, 11, 31^*. This notion is supported by rare cases of individuals with low or no *CETP* expression, who are overall healthy, consequently demonstrating that CETP is not essential for living long and healthy lives*^32–34^*. Lastly, epidemiological studies investigating high mid-life LDL and total cholesterol serum levels showed an association with a greater risk of dementia up to three decades before symptom onset*^35, 36^*. Considering that CETP raises LDL-cholesterol in plasma, its inhibition will reduce this established AD risk factor. Thus, repurposing of safe CETP inhibitors as a therapeutic option for AD holds significant promise, especially in individuals at risk who are still asymptomatic.

Studying the effects of CETP on AD pathology using mice presents a challenge, as mice do not naturally express CETP or homologous proteins*^31, 37, 38^*. Consequently, their lipoprotein profile differs from that of humans. Mouse plasma predominantly contains HDL particles, with lower levels of LDL and very low-density lipoprotein (VLDL) particles, unlike humans who have a higher proportion of LDL and VLDL*^39^*. To address this challenge, a mouse strain expressing the human *CETP* gene under its own promoter was previously generated (CETPtg), resulting in a human-like lipid profile distinct from wild-type (WT) mice and has since been widely used in cardiovascular research*^37^*. In this study, we crossed an AD mouse model of amyloidosis, the McGill-APP-Thy1 (APPtg) mice*^40^* which display age-associated memory deficits, with CETPtg mice and administered the selective CETP inhibitor evacetrapib to investigate the role of CETP in AD as well as determine whether CETP inhibition preserves memory in double transgenic mice (APPtg/CETPtg). Our findings confirm that CETP inhibition preserves memory function in both CETPtg and APPtg/CETPtg mice, suppressing the negative effects of APP. We attributed this maintenance to an altered plasma lipoprotein landscape, differences in brain lipid distribution, and vascular changes in the brain. Most importantly, results from our mouse model were replicated in the human PREVENT-AD cohort consisting of asymptomatic individuals at high risk of developing AD due to a parental history, establishing CETP as a powerful target in AD prevention.

## Methods

### Animal housing

APPtg mice express a human APP-751 transgene carrying the double Swedish (K670N/M671L) and Indiana (V717L) mutations under the Thy1 promoter in neurons on a C57BL/6J background and were kindly provided by Dr. Claudio Cuello*^40^*. The CETPtg strain was purchased from Jackson Laboratory (Jackson Laboratory, Strain #003904) on a C57BL/6J background*^37^*. Husbandry for both strains was heterozygous, and mice were genotyped using Transnetyx. Both strains were crossed to produce CETPtg, APPtg, and double transgenic APPtg/CETPtg mice with WT littermate controls. Heterozygous mice of both sexes were used in this study. Mice were housed in the McGill University Goodman Cancer animal facility. The protocol was approved by the Animal Compliance Office at McGill University (ACO approval #2013-7359).

### Drug preparation

Evacetrapib (LY2484595) was purchased from Selleckchem (Selleckchem, USA). The drug was dissolved in one part Kolliphor EL (Sigma-Aldrich, USA) to stabilize the emulsion of the non-polar compound, and then four parts of 50 g/L D-(+)-glucose (Sigma-Aldrich, USA) were added. The drug solution was kept at 4°C until use.

### Animal exposure and sample collection

Mice were fed a standard chow from Teklad Global Diets T.2920X (Envigo, USA). The diet was changed at 11 weeks of age for the modified low-fat RD Western diet with 1% (w/w) cholesterol (#D16121201 without choline supplementation, Research Diets, USA) until the end of the trial. Mice were randomly divided into the vehicle group (Kolliphor EL-glucose) and the evacetrapib-treatment group receiving daily intraperitoneal injections at 30 mg/kg for 11 weeks*^41^*. Each group comprised 6-8 mice per sex and genotype. Mice had continuous access to food and water. Blood collection (4-6 μL using a capillary tube) was performed at week 12 to determine CETP inhibition in the plasma. Post-mortem, blood was collected using cardiac puncture 4 h after the last injection, plasma was extracted and frozen. For non-perfused mice, the cerebrum was split into four quadrants and frozen in liquid nitrogen. Two lobes of the liver were collected and frozen in liquid nitrogen. Tissues were kept at −80°C. For phosphate-buffered saline (PBS)-perfused mice, left brain hemispheres were fixed in ice-cold 4 % paraformaldehyde (PFA) solutions (Sigma-Aldrich, USA) for 48 h followed by storage at 4°C in sucrose cryoprotective solution until sectioned.

### CETP activity assay

Mouse plasma CETP activity was measured using the CETP activity assay kit (Sigma-Aldrich, USA) using a modified version of the manufacturer’s instructions optimized to a 6-fold reduced volume in a half-area microplate (Greiner Bio-One, Austria). Fluorescence intensity was measured (λ_ex_ = 465/ λ_em_ = 535 nm) using the BioTek Cytation 5 Cell Imaging Multimode Reader (Agilent Technologies, USA).

### Plasma lipid analysis

Mouse plasma analysis was conducted by the Diagnostic Laboratory of Comparative Medicine at the Animal Resources Center at McGill University using a panel for cholesterol, triglycerides, and HDL. The Friedewald equation was used to estimate LDL-cholesterol from HDL-cholesterol and triglycerides*^42^*. VLDL cholesterol was calculated using the Friedewald formula followed by the equation: VLDL cholesterol = triglycerides/2.2mmol/l*^43^*.

### Novel object recognition (NOR) test

A modified version of NOR test based on Ennaceur et al. was used at 21 weeks of age*^44^*. Mice were habituated to handling 3 days prior to the experiment. The experiment was carried out in a sound-proof dark room in an open field arena and with an infra-red video camera recording mouse behaviour. On day one, each mouse was habituated to the box for five minutes. On day two, the familiarization phase, each mouse was put into the same box with two identical objects placed at opposite ends of the box at the equal distance from the nearest corner. Mice were allowed to freely explore the two objects for five minutes. On day three, two objects were placed in the box, one identical to the familiarization phase and the other object was replaced by a novel object. Mice were tested for five minutes while being recorded. Time spent on each object was quantified using the ODLog (Macropod Software, Australia). Mice that spent less than one second on each object were excluded from the analysis. Objects in this study were previously tested for bias. Equipment was cleaned with 70% ethanol and dried between each test to avoid olfactory bias. The novel object discrimination ratio was calculated as follows:

Novel object discrimination ratio (%) = time exploring the novel object/total time spent on both objects *100

### Filipin III staining

Fixed mouse brains were sagittally sectioned at 25 μm thickness using a cryostat (Leica, Germany). The Filipin III staining followed a previous protocol (Oestereich et al.)*^45^*. Sections were mounted on Fisherbrand^TM^ SuperFrost^TM^ (ThermoFisher Scientific, USA) slides using Aqueous Fluoroshield Mounting Medium with DAPI (Abcam, UK). Images were aquired using the widefield Zeiss Axio Observer Fully Automated Inverted Microscope with objective 20X and the ZENPro software v.3.0 (Zeiss, Germany). Images were analyzed using Fiji (ImageJ, USA).

### Aβ ELISA

Human Aβ40 and Aβ42 were analyzed in the mice using an electrochemiluminescence multiplex ELISA using the 6E10 antibody following the manufacturer’s instructions (Meso Scale Discovery, USA). Soluble Aβ was extracted from brain using dimethylamine followed by ultracentrifugation. Insoluble fractions were then extracted from the pellet using formic acid followed by ultracentrifugation to remove debris and neutralized prior to loading onto the plate. Plasma samples were diluted 4-fold. The ELISA was read using the MESO Quickplex SQ 1300 (Meso Scale Discovery, USA).

### Sterol quantification by gas chromatography-mass spectrometry (GC-MS)

Brains and livers were dried in a speedvac concentrator (12 mbar, Thermo Scientific, Germany) and weighed. Sterols were extracted using chloroform/methanol (2:1; v:v) followed by alkaline hydrolysis. Concentrations of sialylated sterols were measured following previous methods*^46^*.

The non-cholesterol sterols (epicoprostanol, ISTD) and the oxysterols (^2^H_x_-oxysterols, ISTD) were detected by mass spectrometry-selected ion monitoring (MS-SIM). GC separation and detection of cholesterol, cholestanol, and 5α-cholestane (ISTD) were performed on a DB-XLB column with a 30 m × 0.25 mm i.d. × 0.25 µm film (J&W Scientific Alltech, USA) in a Hewlett-Packard (HP) 6890 series GC system (Agilent Technologies, USA), equipped with a flame ionisation detector (FID) using 5α-cholestane as an internal standard (ISTD). Non-cholesterol sterols and authentic and deuterium-labelled oxysterols were separated on a second DB-XLB column in an HP 6890N Network GC system connected with a direct capillary inlet system to an inert quadruple mass selective detector HP5975B (Agilent Technologies, Germany). Several internal controls for the plasma or tissue extract were used (Supplemental Table 1). To avoid autoxidation, 50 µL of a 2.6.-di-tert.-butylmethylphenol/methanol solution (Sigma-Aldrich, USA) were added. After saponification with 2 mL of 1 M 95% ethanolic sodium hydroxide solution (Merck, Germany) (60°C, 1 h) free sterols and oxysterols were extracted three times with 3 mL of cyclohexane each. Organic solvent was evaporated by nitrogen stream (60°C). Residues were dissolved in 80 µL n-decane (Merck, Germany). An aliquot (40 µL) was incubated (70° C, 1h) in addition to 20 µL of trimethylsilylating (TMSi) reagent-comprising chlortrimethylsilane (Merck, Germany), 1.1.1.3.3.3-hexamethyldisilasane (Sigma-Aldrich, USA), and pyridine (Merck, Germany, 9:3:1) in a GC vial for GC–mass selective detector non-cholesterol and oxysterol analysis. Another aliquot (40 µL) was incubated in addition to 40 µL of the TMSi-reagent and dilution with 300 µL of n-decane in a GC vial for GC–FID cholesterol analysis. 2 µL were injected by automated injection in splitless mode using helium (1 mL/min) for GC–MS-SIM and hydrogen (1mL/min) for GC–FID analysis (injection temperature 280°C). The temperature was kept at 150°C for 3 min, then heated at 20°C/min, and kept at 290°C for 34 min. For mass selective detection, electron-impact ionisation was applied with 70 eV. MS-SIM was performed by cycling the quadruple mass filter between different m/z values at a rate of 3.7 cycles/s. TMSi derivatives of the non-cholesterol sterols and di-TMSi-derivatives of the oxysterols monitoring compounds and specific m/z were listed (Supplemental Table 2).

Peak integration was performed manually. Cholesterol and cholestanol were directly quantified by multiplying the ratios of the area under the curve (AUC) of cholesterol or cholestanol to 5α-cholestane by 50 µg (ISTD amount). Non-cholesterol sterols and oxysterols were quantified by comparing the AUC ratios of each to internal standards, using MS-SIM analyses and standard curves for the respective sterols/oxysterols. Identification of all sterols was verified by comparison with the full-scan mass spectra of authentic compounds. Additional qualifiers (characteristic fragment ions) were used for structural identification (m/z values not shown).

### RNA extraction and RNA-sequencing analysis

Total RNA was extracted from bulk brain tissue of CETPtg and APPtg/CETPtg mice using the Qiagen RNeasy Plus Universal Mini Kit following the manufacturer instructions (Qiagen, Germany). RNA integrity number (RIN) factor was measured for samples ranging between 7.3 and 8.6 (Agilent Technologies, USA). mRNA was enriched from 100-500 ng total RNA using NEBNext Poly(A) mRNA Magnetic Isolation Module Kit, and libraries were generated using NEBNext Ultra Directional RNA library kit (New England Biolabs, USA). These cDNA libraries were sequenced using the Illumina NovaSeq X 25B sequencer (Illumina, USA), 150 nucleotide paired-end reads. RNA-seq fastq reads were trimmed using Trimmomatic (version 0.39) to remove low-quality bases and adapters. Trimmed reads were aligned to the *mus musculus* 10 (mm10) reference genome using STAR (version 2.7.8a) with default parameters to generate .bam files. Alignments were sorted and duplicates were marked using Picard (version 2.0.1). Raw and normalized reads were quantified using HTSeq-count and the StringTie (version 1.3.5) respectively. Differentially expressed genes (DEGs) with a maximum adjusted p-value of 0.05 and at least 1.05 absolute value fold change were identified using DEseq2 (Bioconductor release 3.19). Samples were hierarchically clustered according to their gene expression profiles across the DEGs. Gene expression was normalized by calculating the z-scores of FPKMs within each gene. Clustering was performed by Euclidean distance using Ward’s method.

Gene ontology (GO) terms significantly enriched in the DEGs were determined using clusterProfiler (Bioconductor release 3.19) with a maximum Benjamini-Hochberg adjusted p-value (p.adj) of 0.05 with the biological process, cellular component, and molecular function gene sets. Redundant GO-terms were removed using the simplify function in clusterProfiler. DEGs were analyzed for enrichment in GO ontologies for biological process, cellular component, and molecular function. Gene set enrichment analysis (GSEA) (version 4.3.3) was performed using the normalized read counts produced from DESeq2*^47^*. The analysis used default parameters, including 1000 permutations, and the biological process gene sets in the Molecular Signatures Database (MSigDB), showing significant enrichment in the GOBP_BLOOD_VESSEL_REMODELING and GOBP_REGULATION_OF_ENDOTHELIAL_CELL_DIFFERENTIATION sets.

### Human PREVENT-AD Cohort

The PREVENT-AD longitudinal cohort is composed of cognitively unimpaired individuals who have a first degree relative with AD and therefore are at a higher risk of developing this dementia*^48^*. The majority of participants were over the age of 60 at recruitement. However, individuals aged 55-59 years were included if they were within 15 years of the onset of their youngest-affected relative’s symptoms. Over 370 participants have been monitored annually for the past 13 years with biochemical, clinical and cognitive assessments, cerebrospinal fluid (CSF) and blood biomarkers and neuroimaging modalities (structural and functional magnetic resonance imaging (MRI), and amyloid and tau positron emission tomography (PET) scans)*^48^*. Data used in the preparation of this article were obtained from the PREVENT-AD program (https://douglas.research.mcgill.ca/stop-ad-centre), data release 7.0. A complete listing of the PREVENT-AD Research Group can be found in the PREVENT-AD database (https://preventad.loris.ca/acknowledgements/acknowledgements.php?date=2024-10-01.)

### CSF Aß42, total tau (t-tau), and phosphorylated tau (p-tau181) analysis

Lumbar punctures were performed in a subset of volunteers (n=120) in the morning following an overnight fast, with a Sprotte 24-gauge atraumatic needle as described in Tremblay-Mercier et al (2021)*^49^*. CSF was centrifuged for 10 minutes at room temperature, cells and insoluble material were excluded, and aliquots were stored at –80°C. AD biomarkers (p-tau181, t-tau, and Aß42) were measured according to procedures developed by the BIOMARKAPD consortium, using validated Innotest enzyme-linked immunosorbent assay kits (Fujirebio, Japan)*^50^*.

### The Repeatable Battery for the Assessment of Neuropsychological Status (RBANS)

In the PREVENT-AD cohort, cognitive status is assessed annually by RBANS scale. RBANS was developed with the purpose of detecting and characterizing early dementia in older adults as well as in younger individuals*^51^*. RBANS tests assess five cognitive domains: immediate memory, visuospatial/constructional skills, attention, language, and delayed memory. Scores are calculated individually for each cognitive domain and then combined to provide a total score. The development and validation of the RBANS is described in detail by Randolph et al (1998)*^51^*.

### Analysis of human CSF proteomic profile by SOMAscan

The SOMAscan panel measured ~7500 aptamers mapping to approximately 6600 unique protein targets. Protein measurements are reported in relative fluorescence unit (RFU). Initial data normalization procedures were performed by SomaLogic. Briefly, hybridization normalization was performed at the sample level. Aptamers were then divided into three normalization groups: S1, S2 and S3; based on the observed signal to noise ratio in technical replicates and samples. This division was done to avoid combining features with different level of protein signal for additional normalization steps. Median normalization was then performed to remove other assay biases such as protein concentration, pipetting variation, variation in reagent concentrations, and assay timing among others*^52^*. Finally, normalization to a reference was performed on individual samples to account for additional technical variance as well as biological variance. This normalization step was performed using iterative Adaptive Normalization by Maximum Likelihood (ANML), a modification of median normalization, until convergence is reached. Additional details on normalization procedures are documented in SomaLogic’s technical note*^53^*.

### Statistical analyses for the mouse dataset

Statistical analyses were run using the GraphPad Prism software version 9, where statistical significance was considered *p* < 0.05. A two-way ANOVA was used to evaluate the effects of treatment and genotype in biochemical and behavioural analyses. When significant main effects or interactions were identified, Tukey’s multiple comparison post-hoc tests were conducted.

### Statistical analyses for the human SOMAscan dataset

Relationships between CSF protein levels at participants’ baseline visit were analyzed using general linear models (GLMs), adjusting for participant age, *APOE* ε4 status, and sex. Further GLMs were performed to test the interaction between sex and baseline protein levels on the dependent variable. To assess sex-specific effects, analyses were then stratified by sex using simple effects tests, regardless of the interaction term’s statistical significance.

The trajectory of participants’ cognitive status by RBANS was determined by calculating the slope of scores from their baseline visit to their most recent assessment within a seven-year period. GLMs were run to investigate the effect CETP or AD markers (p-tau181, t-tau, Aβ42) in the CSF at baseline on the trajectory of cognition, adjusting for participant age, *APOE* ε4 status, sex, and years of education. GLMs testing the interaction between sex and baseline CETP or AD marker levels in the CSF were performed, followed by sex-stratified simple effects analyses. To assess if baseline CETP and/or sex moderates the effect of baseline CSF p-tau181/Aβ42 levels on cognitive trajectory, additional GLMs were run testing this three-way interaction (Baseline CSF Aβ42 levels x sex x high/low baseline CSF CETP levels), proceeded by simple effects analyses stratified by sex and CETP grouping (regardless of the interaction term’s statistical significance).

Statistical analyses were run using the Jamovi software version 2.5, where statistical significance was considered *p* < 0.05*^54–58^*. For all GLM analyses, if the model’s residuals did not meet the assumption of normal distribution, data were sinh-arcsinh (SHASH) transformed using JMP® Pro version 17.0.0. Robust standard errors (HC3) were used for analyses when the homoskedasticity assumption was not met (Supplemental tables 3, 4).

## Results

### Evacetrapib successfully inhibits CETP in mice

To explore the impact of CETP on AD pathophysiology in a preclinical model, APPtg mice and CETPtg mice were crossed to generate double transgenic APPtg/CETPtg. APPtg, CETPtg and WT mice were used as littermate controls (Fig. 1A). Mice were fed a regular chow from birth until week 11, and then switched to a 1% cholesterol diet based on Oestereich et al. to induce CETP expression and increase plasma LDL levels in CETP-expressing mice*^45^*. Alongside the 1% cholesterol diet, mice received either vehicle or evacetrapib at 30 mg/kg/day intraperitoneally for 11 weeks (Fig. 1B). To assess drug-target engagement in our model, we measured CETP enzymatic activity in plasma during the trial and at endpoint (Fig. 1C-H). CETP activity was significantly reduced 4 h after evacetrapib treatment compared to the vehicle group (Fig. 1D). At 24 h, CETP activity rebound while still being reduced showing a transient inhibition of our dosage (Fig. 1E). As expected, no CETP activity was detected in WT and APPtg mice. Similarly, terminal plasma collection 4 h after evacetrapib administration showed that CETPtg and APPtg/CETPtg mice exhibited significantly lower CETP activity than their vehicle-treated counterparts (Fig. 1F-H), confirming effective drug-target engagement in our mouse model.

**Figure 1:**
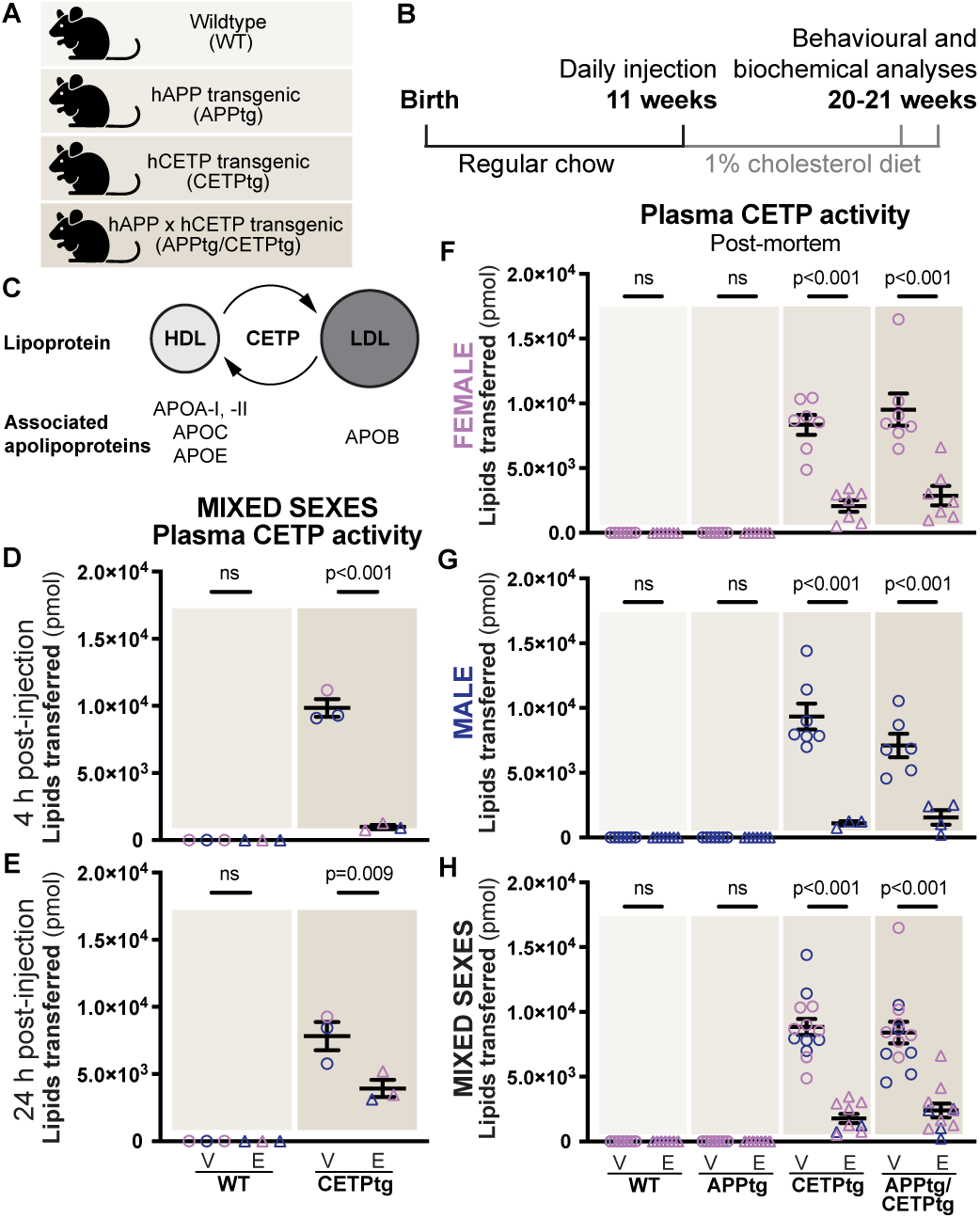
Evacetrapib treatment decreases CETP activity. **A** Scheme of the experimental design created with BioRender.com. **B** Timeline of the mouse study including behavioural assays, and organ collection. **C** Scheme of how plasma CETP transfers neutral lipids between ApoA-I-containing HDL and ApoB-containing LDL particles. **D, E** CETP activity quantification at 4 h or 24 h post-injection after two weeks of treatment in a subset of mice. **F-H** CETP activity measured at the time of sacrifice in female and male mice, *n*=*6-8,* as well as both sexes combined, *n=12-14*. **D-H** Data is shown as bar charts indicating mean ± SEM with individual points. Circles represent the vehicle (V) condition while triangles denote the evacetrapib (E) condition. Female data points are represented in violet and male data points in indigo. Statistical significance was measured using two-way ANOVA followed by Tukey’s post-hoc test.

### Evacetrapib remodels the lipoprotein profile in mice

As a result of CETP activity and inhibition, the plasma lipoprotein profile is expected to change. As in previous reports*^37, 38^*, CETP activity increased LDL and decreased HDL levels in vehicle-treated CETP-expressing mice as compared to WT or APPtg mice (Fig. 2A-F). However, such lipoprotein changes clearly occurred in female mice and were attenuated in male mice (Fig. 2A-D). Accordingly, CETP inhibition in female mice normalized LDL levels back to baseline, while HDL levels remained higher than baseline levels compared to WT mice on vehicle (Fig. 2A, B). When data from both sexes were pooled, significant differences in HDL and LDL were mostly lost, highlighting the importance of sex stratification in this analysis (Fig. 2E, F). A correlation analysis showed a positive correlation between CETP activity and LDL, and a negative correlation with HDL except for HDL in males (Suppl. Fig. 1A-F). Total plasma cholesterol, triglyceride and VLDL levels remained consistent across conditions and genotypes in both male and female mice (Suppl. Fig. 1G-O).

**Figure 2:**
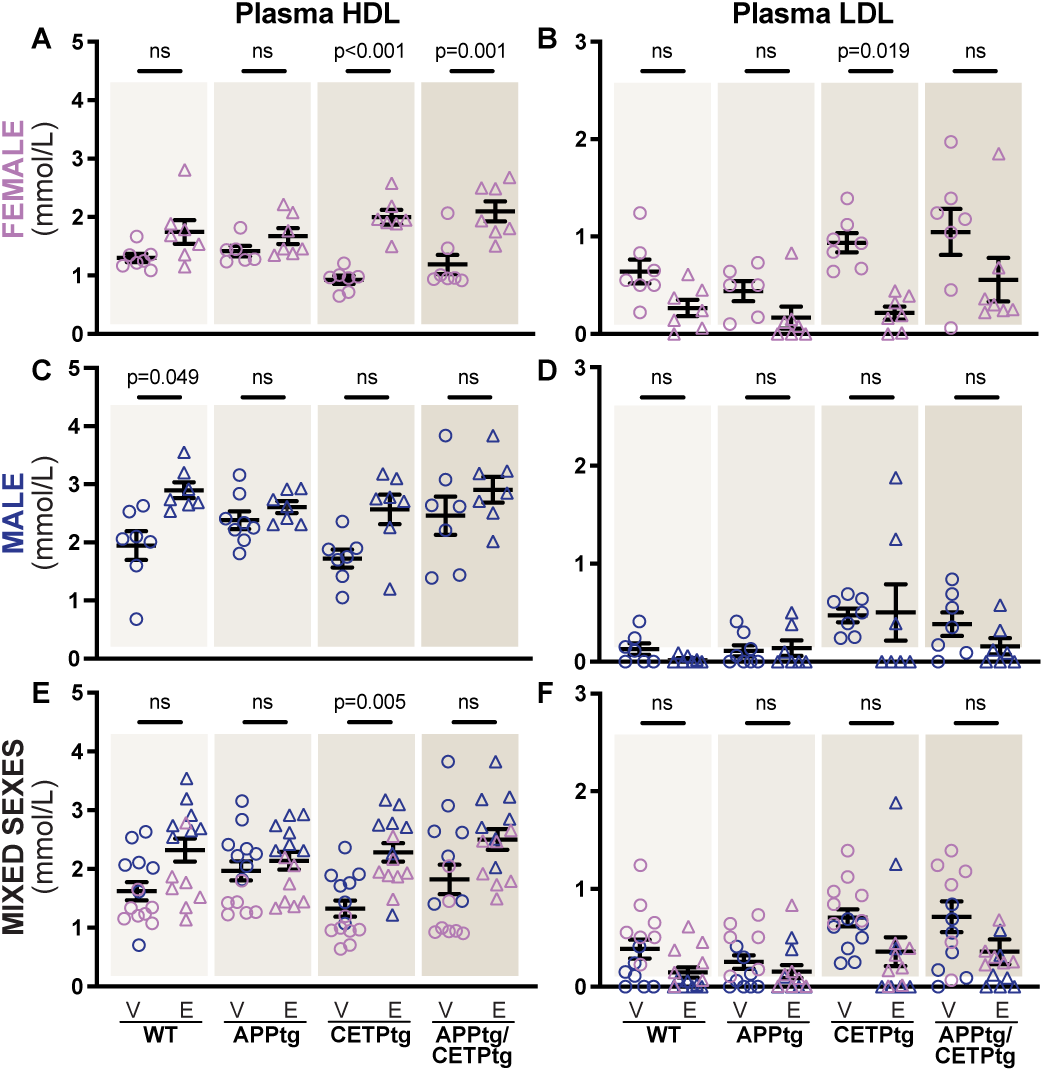
Evacetrapib treatment changes the lipoprotein profile in CETP expressing mice. **A-F** Plasma lipoproteins HDL and LDL were quantified in female and male mice, *n*=*6-8,* as well as both sexes combined, *n=12-14*. Data is shown as bar charts with individual points. Circles represent the vehicle (V) condition while triangles denote the evacetrapib (E) condition. Bar symbols indicate mean ± SEM, where female data points are represented in violet and male data points in indigo. Statistical significance was measured using two-way ANOVA followed by Tukey’s post-hoc test.

### Evacetrapib maintains memory in APPtg/CETPtg mice

To reveal whether CETP activity affects cognitive performance in our new mouse model similarly to previously described human epidemiological studies, we assessed recognition memory using the NOR test. Mice were assessed at 21 weeks of age, which is the earliest stage at which cognitive impairment becomes detectable in APPtg mice (Fig. 3A-D). WT mice showed no impairment, regardless of sex or treatment, demonstrating that evacetrapib had no negative impact on memory. APPtg mice, both male and female, displayed expected memory impairment, independent of treatment, showing that APP abundance perturbs memory through a mechanism that is not affected by evacetrapib. Interestingly, CETP expression alone induced memory impairment in female mice (Fig. 3C). However, memory was sustained with evacetrapib, demonstrating a pharmacological way to prevent memory loss. The double transgenic APPtg/CETPtg female mice exhibited memory impairment as well, which could be due to the presence of APP or CETP since both impaired memory functions alone. However, we cannot infer any potential additive effects. Strikingly, evacetrapib treatment preserved memory function in APPtg/CETPtg female mice despite APP abundance (Fig. 3C) showing that CETP inhibition dominates over the APP-mediated memory impairment. This result establishes CETP inhibition as a viable strategy to delay memory impairment. Males did not show significant differences, though similar trends can be observed, between evacetrapib- and vehicle-treated CETP expressing groups. Therefore, further analyses were continued in female mice (and results from male mice are presented in the supplemental data). We further assessed memory performance using the Y-maze test which evaluates spatial memory. No changes, including no impairment in APPtg mice, were observed indicating that spatial memory remains unaffected at this early stage (Suppl. Fig. 2A-C)*^59^*.

**Figure 3:**
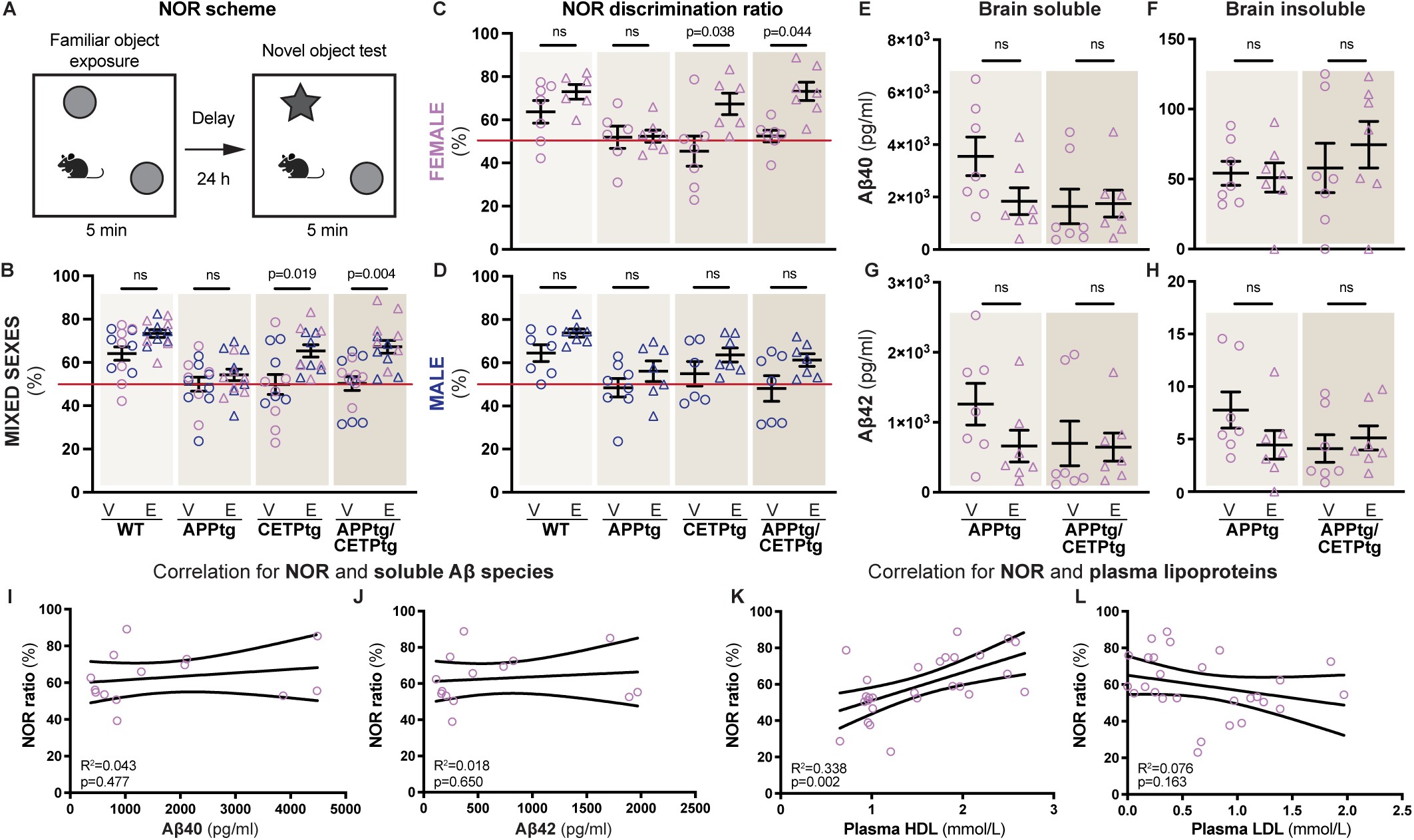
Memory in CETP-expressing mice is preserved with evacetrapib. **A** Scheme of the NOR test created with BioRender.com. **B-D** NOR scores are displayed for females, males, *n*=*6-8,* and both sexes combined, *n=12-14*, with a red line indicating the 50% discrimination ratio at which mice cannot differentiate between the familiar and the novel object suggesting memory impairment. **E-H** Soluble and insoluble Aβ40 and Aβ42 peptides were quantified in brain homogenates of female mice by ELISA. **B-H** Data is shown as bar charts with individual points. Circles represent the vehicle condition while triangles denote the evacetrapib condition. Bar symbols indicate mean ± SEM, *n*=*6-8,* where female data points are represented in violet and male data points in indigo. Statistical significance was measured using two-way ANOVA followed by Tukey’s post-hoc test. **I-L** Linear correlations of Aβ40 and Aβ42 versus cognition (NOR), *n*=*14*, (**I-J)** as well as HDL and LDL versus cognition (NOR), *n*=*27-28* (**K-L)** in female APPtg and APPtg/CETPtg mice on evacetrapib and vehicle control. Data is shown as scatter plots with a simple linear regression line. Correlations were analyzed using Pearson’s correlation assuming normality and the 95% confidence interval is shown on the plot as dotted lines. R-squared (R^2^) and p-values (p) are indicated for each correlation.

### CETP transgenicity does not exacerbate the amyloidogenic pathway

According to the amyloid hypothesis, APPtg mice develop cognitive impairment due to the generation of Aβ peptides by proteolytic cleavages of APP*^60^*. To reveal whether CETP activity impacts Aβ production, we quantified Aβ40 and Aβ42 peptides from brain lysates at 21 weeks of age by multiplex ELISA. No significant differences were observed in either soluble or insoluble (plaque deposited) Aβ species across conditions (Fig. 3E-H). Similar results were found in males (Suppl. Fig. 3A-D). Thus, we infer that the memory rescue by evacetrapib in APPtg/CETPtg is not due to altered Aβ peptide production. Further, we quantified Aβ peptides from female mouse plasma and found no changes between groups indicating that evacetrapib did not change the Aβ clearance rate in our mouse model (Suppl. Fig. 3E, F). Accordingly, neither plasma to brain Aβ ratios nor brain Aβ42/Aβ40 ratios showed differences (Suppl. Fig. 3G-J). Likewise, correlation analyses did not find interactions between memory and Aβ in our model (Fig. 3I, J). Overall, the absence of significant differences in Aβ production or clearance between evacetrapib- and vehicle-treated mice in the APPtg/CETPtg group shows that evacetrapib preserves memory independent of the amyloidogenic pathway.

### Good memory correlates with high HDL levels

Previous studies have shown that high HDL levels protect from memory decline*^61^*. Consequently, we correlated memory performance with lipoprotein levels. We found a significant positive correlation of good memory performance with HDL levels in female mice (Fig. 3K). LDL levels showed a negative trend with cognitive performance, though not significant (Fig. 3L). Thus, one explanation for the maintained for memory in females could be the high HDL levels.

### CETP expression does not increase cerebral cholesterol synthesis or excretion

The majority of brain cholesterol is synthesized locally by oligodendrocytes and astrocytes*^62–64^*. Therefore, cerebral CETP activity may affect local brain cholesterol synthesis independently from plasma cholesterol. We tested the possibility that CETP affects either 1) cholesterol synthesis, 2) cholesterol excretion, or 3) cholesterol distribution in the brain (Fig. 4A). Using MS, we measured levels of cholesterol precursors, cholesterol derivatives, and total cholesterol in bulk brain tissue (Fig. 4B-J). First, we examined the abundance of cholesterol precursors lanosterol and desmosterol from the Bloch pathway in female brains and noted no variations between the groups (Fig. 4B-D). Similar results were observed for cholesterol precursors in the Kandutsch-Russell pathway (Fig. 4E-G). The precursor ratios demonstrate that there are no blockages in the respective pathways (Fig. 4D, G). Next, we measured the amount of total cholesterol in the brains of female mice and with no changes reported (Fig. 4H). Further, previous studies have shown that defects in prenylation, a process related to cholesterol synthesis, impacts memory*^65, 66^*. We investigated mRNA and protein amounts for enzymes involved in the mevalonate/isoprenoid pathway, specifically farnesyl diphosphate synthase (FDPS), farnesyl-diphosphate farnesyltransferase 1 (FDFT1), and geranylgeranyl diphosphate synthase 1 (GGPS1), and found no differences between evacetrapib and vehicle-treated mice, therefore excluding their involvement in memory maintenance in our model (Suppl. Fig. 4A-G). We conclude that cholesterol synthesis is not affected by CETP activity in our model. To assess cholesterol excretion, cholesterol derivatives 24S-hydroxycholesterol and 27-hydroxycholesterol were quantified in both brain and plasma, as well as brain cholestanol. However, no differences were found, excluding an effect of CETP on cholesterol excretion (Fig. 4I, J, Suppl. Fig. 5A). BBB integrity was indirectly assessed by measuring diet-derived plant sterols both in brain and plasma, which showed no changes indicating that the BBB had comparable functionality between groups (Suppl. Figs. 5B-D, 6A-C). Further, no significant differences were found in plasma and liver samples for cholesterol, cholesterol metabolites, or cholesterol precursors (Suppl. Figs. 6A-Z, 7A-AF). Thus, we concluded that the memory preservation associated with CETP inhibition does not derive from altered cholesterol synthesis or excretion.

**Figure 4:**
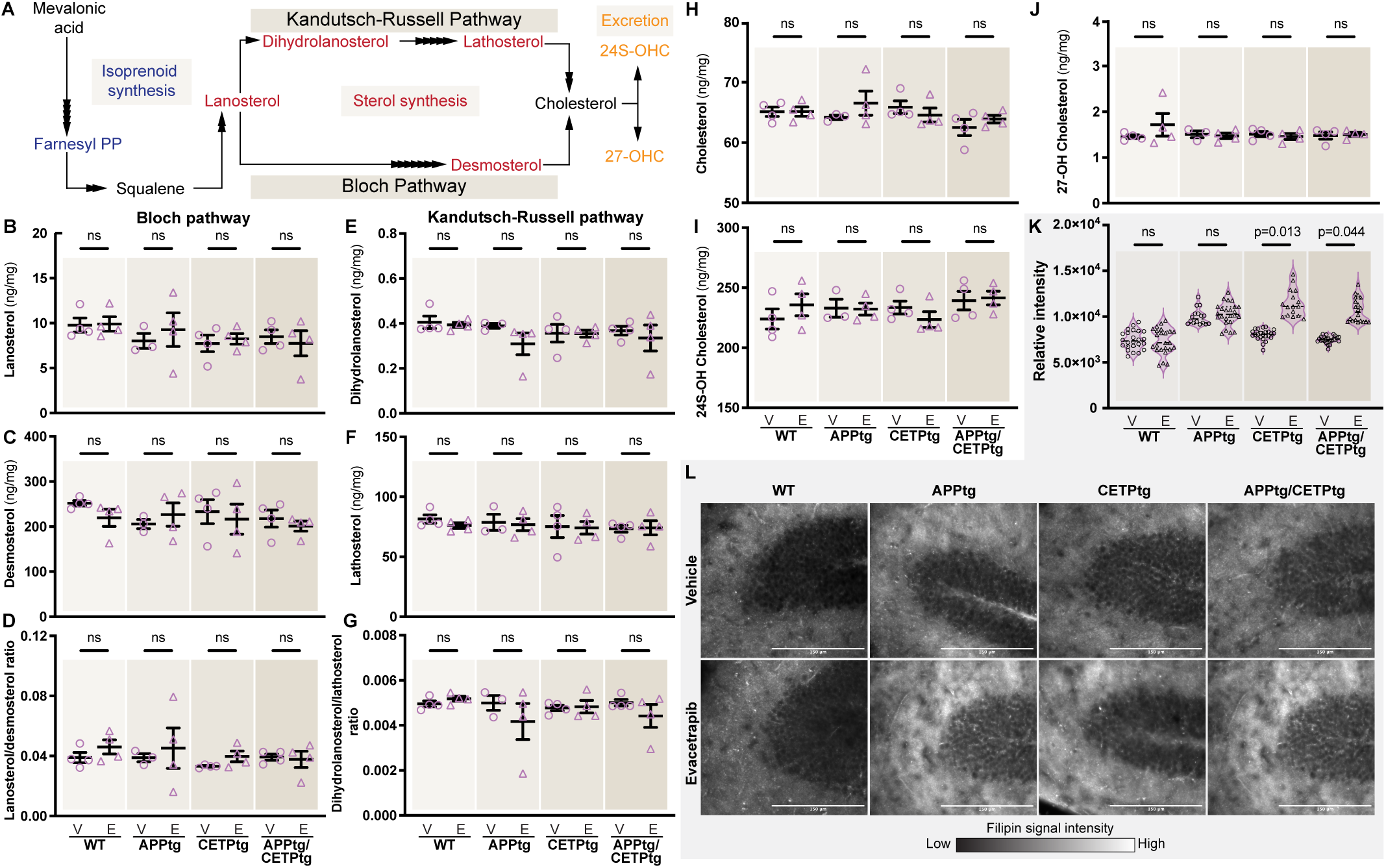
Evacetrapib treatment changes lipid distribution in the hippocampus without affecting overall sterols, sterol precursors or sterol derivatives in the brain. **A** Scheme of the mevalonate pathway, highlighting key steps of isoprenoid and cholesterol synthesis with cholesterol metabolites. **B, C, E, F** Lanosterol and desmosterol were detected from brain tissue, along with dehdrolanosterol and lathosterol. **D, G** Ratios of lanosterol to desmosterol, and 24.25-dihydrolanosterol to lathosterol. **H-J** Total brain cholesterol was quantified along with cholesterol derivatives 24(S)-hydroxycholesterol (24(S)-OH) and 27-hydroxycholesterol (27-OH). **B-J** Data is shown as bar charts indicating mean ± SEM with individual points, *n=3-4*, where female data points are represented in violet. **K, L** Filipin III staining in the dentate dyrus of the hippocampus in female mice was quantified (**K**). Data is shown as violin plots with individual measurements (4-8 technical replicates) from each biological replicate *n=3* represented as a data point with the median (full line), first and third quartile (dotted lines). **L** Representative images of sagital sections of three biological replicates at 40X magnification are shown. Signal intensity is depicted using a scale bar. Scale 150 µm. **B-K** Circles represent the vehicle condition while triangles denote the evacetrapib condition. Statistical significance for all analyses was measured using two-way ANOVA followed by Tukey’s post-hoc test.

### CETP expression changes the lipid distribution in the hippocampus of CETP expressing mice

To complement the bulk biochemical cholesterol analyses, we next examined the spatial distribution of cholesterol and neutral lipids in brain sections. To detect free cholesterol, we used filipin staining*^67^*. When comparing hippocampal cholesterol content across the different genotypes treated with the vehicle, we observed that APPtg mice exhibited approximately 50% more free cholesterol than WT mice, a finding consistent with previous reports*^68, 69^*. CETPtg and APPtg/CETPtg mice showed an average increase of approximately 10% in free cholesterol compared to WT mice, in line with our previous findings*^45^*. Notably, evacetrapib treatment in CETPtg and APPtg/CETPtg mice further increased total cholesterol levels by up to 60% as compared to WT, specifically in the dentate gyrus (Fig. 4K, L). This result was unexpected as APPtg mice, regardless of treatment, showed higher free cholesterol but exhibited memory impairment. In contrast, evacetrapib-treated CETPtg and APPtg/CETPtg mice maintained their memory despite having higher cholesterol levels. We conclude that the absolute amount of hippocampal cholesterol alone is not determining memory performance. Next, we evaluated the abundance of neutral lipids in the hippocampus using BODIPY665/676 staining, which mostly stains lipid droplets (triglycerides and cholesteryl esters)*^70^*. A trend was observed for CETPtg and APPtg/CETPtg mice on evacetrapib showing lower neutral lipid levels compared to all other groups, though not significant (Suppl. Fig. 8A-B). The ratio of free cholesterol to neutral lipids was elevated in evacetrapib-treated CETPtg and APPtg/CETPtg mice (Suppl. Fig. 8C). Thus, while we detected no lipid changes with bulk GC-MS analysis, the spatial information gained by analyzing brain sections revealed that evacetrapib leads to a significant redistribution of cholesterol and a remodelling of lipid stores, specifically in the hippocampus, providing an additional explanation for the maintenance of memory by CETP inhibition.

### Expression of genes involved in blood vessel remodeling are altered by CETP inhibition

To narrow down the mechanism by which CETP inhibition maintains memory in our mouse model, we performed an unbiased analysis of gene expression comparing evacetrapib- and vehicle-treated groups of CETP-expressing female mice. We identified differentially-expressed genes (DEGs) and found an upregulation of 19 genes and downregulation of 5 genes (fold change ≥ |1.05|, *q*-value ≤ 0.05) (Fig. 5A). Hierarchical clustering within each genotype (CETPtg and APPtg/CETPtg) showed that samples cluster by treatment based on their expression profiles across the DEGs, confirming that the effects of evacetrapib on gene expression are consistent within each genotype (Fig. 5B). Out of the 24 DEGs, 11 (*Tagln, Acta2, Myh11, Pecam1, Flt1, Kdr, Cdh5, Mmrn2, Podxl, Tgm2, Cd59a*) are expressed in cells involved in brain barriers, with six (*Pecam1, Flt1, Cdh5, Mmrn2, Podxl, Cd59a*) specifically expressed in brain endothelial cells, four (*Tagln, Acta2, Myh11, Kdr*) in vascular smooth muscle cells (VSMCs) at the choroid plexus, and one (*Tgm2*) expressed both in endothelial cells and VSMCs*^71–74^*. Furthermore, although five of the 24 genes (*Ppp1r10, Ccnd1, Tnfsf10, Atp2a3, Osmr*) are not specifically expressed in brain barrier cells, they are documented to play important roles in vascular function*^75–79^*. Unbiased GO term analysis revealed a positive enrichment for terms related to blood vessel maintenance and angiogenesis (Fig. 5C, D). In a second approach, we used GSEA analysis to confirm the significant dysregulation of a gene set associated with a particular biological process (Fig. 5E, F). Among the remaining DEGs that are neither expressed in brain barriers nor involved in vascular functions, four genes (*Rims4, Ppp1r9b, Wwc1, Atp2c2*) are associated with neuronal function, memory, and synaptic plasticity, while two genes (*Per2, Herpud1*) are linked to circadian rhythms *^71–73, 80–84^*. A principal component analysis was performed for quality assessment (Suppl. Fig. 9A).

**Figure 5:**
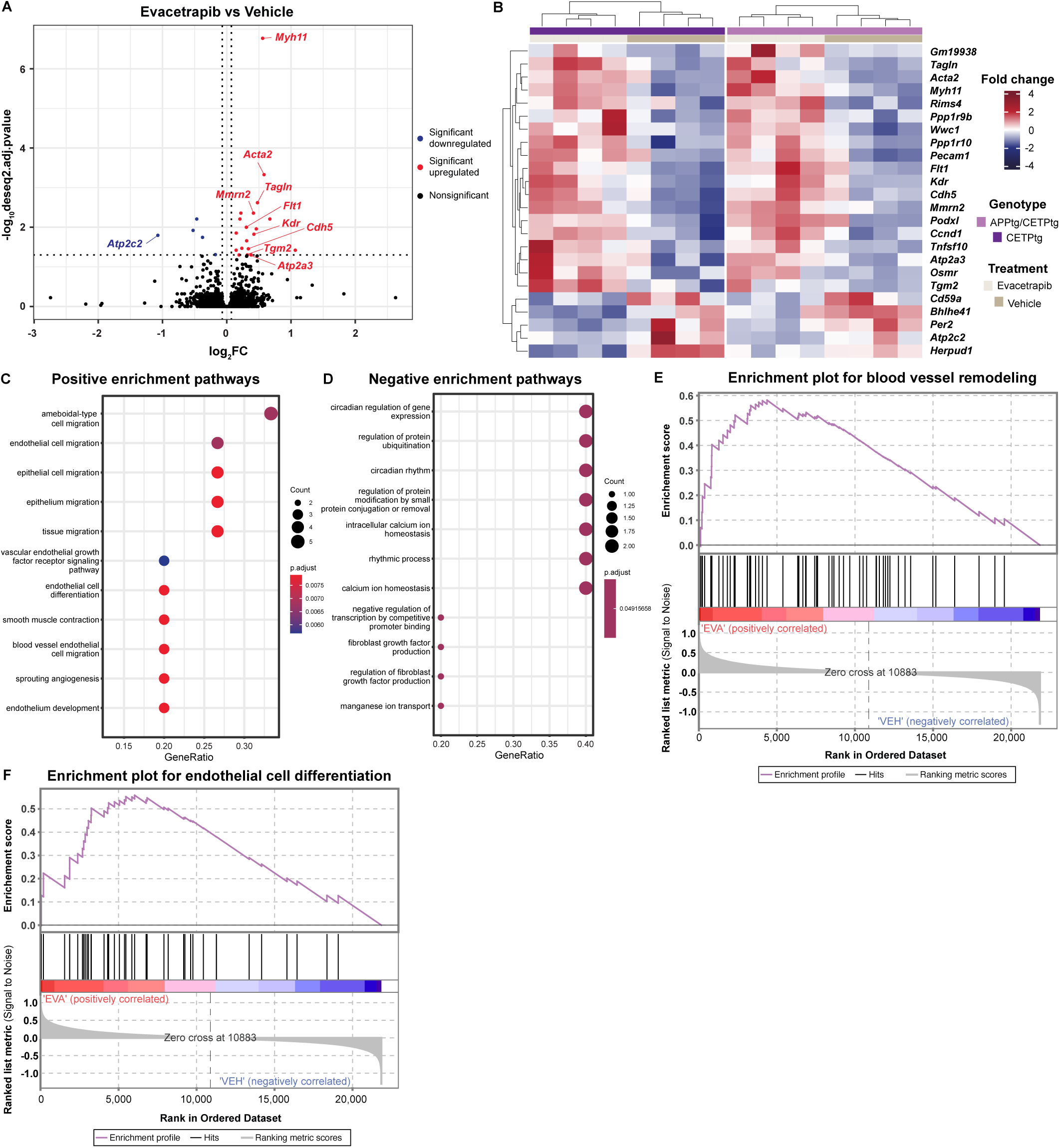
Evacetrapib upregulated genes linked to brain barrier functions. **A** Volcano plot of differential gene expression analysis comparing evacetrapib- and vehicle-treated female mice, *n=8,* by RNAseq. Significant DEGs with |fold change| ≥ 1.05 and *q*-value ≤ 0.05 are shown in red, indicating upregulation, or blue, indicating downregulation. **B** Heatmaps and hierarchical clustering of CETPtg and APPtg/CETPtg female mice treated with evacetrapib or vehicle, *n=4*, based on normalized expression of DEGs. The panel shade approaching red indicates a high expression, while the blue shade indicates low expression. **C, D** Bubble plots showing upregulated and downregulated GO enrichment analysis of DEGs acquired by RNAseq. The size of the circle at each individual GO terms represents the number of genes enriched. The dot shade approaching red indicates higher significance. **E, F** GSEA enrichment plots using the GO-term database were generated for evacetrapib-treated and vehicle-treated CETPtg and APPtg/CETPtg female mice, *n*=*4*. The analysis runs along the ranked list of genes (left to right) with the running enrichment score (violet line) represents the enrichment of the gene set across the ranked list.

### Association of CETP abundance in the CSF with other proteins in the PREVENT-AD cohort

To assess whether the findings of our mouse model reflect disease-changes in humans, we tested if key findings of our mouse model are observed in the PREVENT-AD cohort*^49^*. The PREVENT-AD cohort recruits pre-symptomatic subjects before the appearance of mild cognitive impairment (MCI), but with a strong parental predisposition to developing AD. Over time, some participants (~20%) have progressed to MCI allowing longitudinal cognitive analyses and identifying biochemical changes that occur in the early stages of AD. Cognitive assessment was done with the RBANS over the course of 13 years. The visuospatial constructional index score (VCIS) is one of the RBANS subcategories. Individual trajectories (slopes) of cognitive performance were calculated using cognitive assessment scores from baseline to their most recent follow-up within a set seven-year period. There was a significant difference in the effect of CETP on VCIS trajectory between sexes, where high CETP levels in the CSF at baseline were associated with a decline in VCIS over time in women indicating that low CETP levels, which associates with low CETP activity*^85^*, is related to better cognitive performance like in our mouse model (Fig. 6A). VCIS showed no relationship with Aβ42 alone, suggesting CSF Aβ42 levels do not impact cognition at this stage similarly to APPtg/CETPtg mice (Fig. 6B). However, we observed a significant effect of the CSF p-tau(181)/ Aβ42 ratio, a strong predictor of worsening cognition, on VCIS suggesting conversion to MCI and potentially AD in these individuals (Fig. 6C; Suppl. Fig. 10A). Further regression analyses between RBANS, CETP and AD markers as well as non-differentially expressed proteins are shown in the supplemental figure (Suppl. Fig. 10C-J). A subset of CSF samples underwent SOMAscan proteomic analysis at baseline including CETP protein levels allowing us to examine the effect of CSF CETP levels on the DEG products identified by the RNA sequencing from mouse brain tissue. Out of 24 DEGs, 23 are protein-coding genes (excluding Gm19938), of which SOMAscan has probes for 19 proteins. 14 proteins were successfully quantified in the CSF of PREVENT-AD participants by SOMAscan (TAGLN (also known as transgelin), PPP1R9B (spinophilin), WWC1, PPP1R10, PECAM1 (CD31), FLT1 (vascular endothelial growth factor receptor 1, VEGFR1), KDR (vascular endothelial growth factor receptor 2, VEGFR2), CDH5 (cadherin-5), MMRN2, TNFSF10, ATP2A3, OSMR, TGM2 (transglutaminase-2), CD59) (Fig. 6D-L; Suppl. Fig. 10H-J). Out of the 14 detected proteins, 11 were differentially regulated in relationship to CSF CETP levels in women (nine brain barrier cell-associated proteins, and two neuronal proteins, Fig. 6D-N), showing that changes in our CETP mouse model are relevant in presymptomatic AD humans. The cohort demonstrated no significant change in CSF CETP protein levels as a function of the CSF/plasma albumin ratio or the CSF/serum IgG ratio showing that the BBB was intact in these asymptomatic subjects (Suppl. Fig. 10K, L). We conclude that the majority of DEG identified in the mouse model are found with differential abundance in the CSF in at-risk humans as a function of CETP levels, demonstrating that our mouse model replicates aspects of human disease progression.

**Figure 6:**
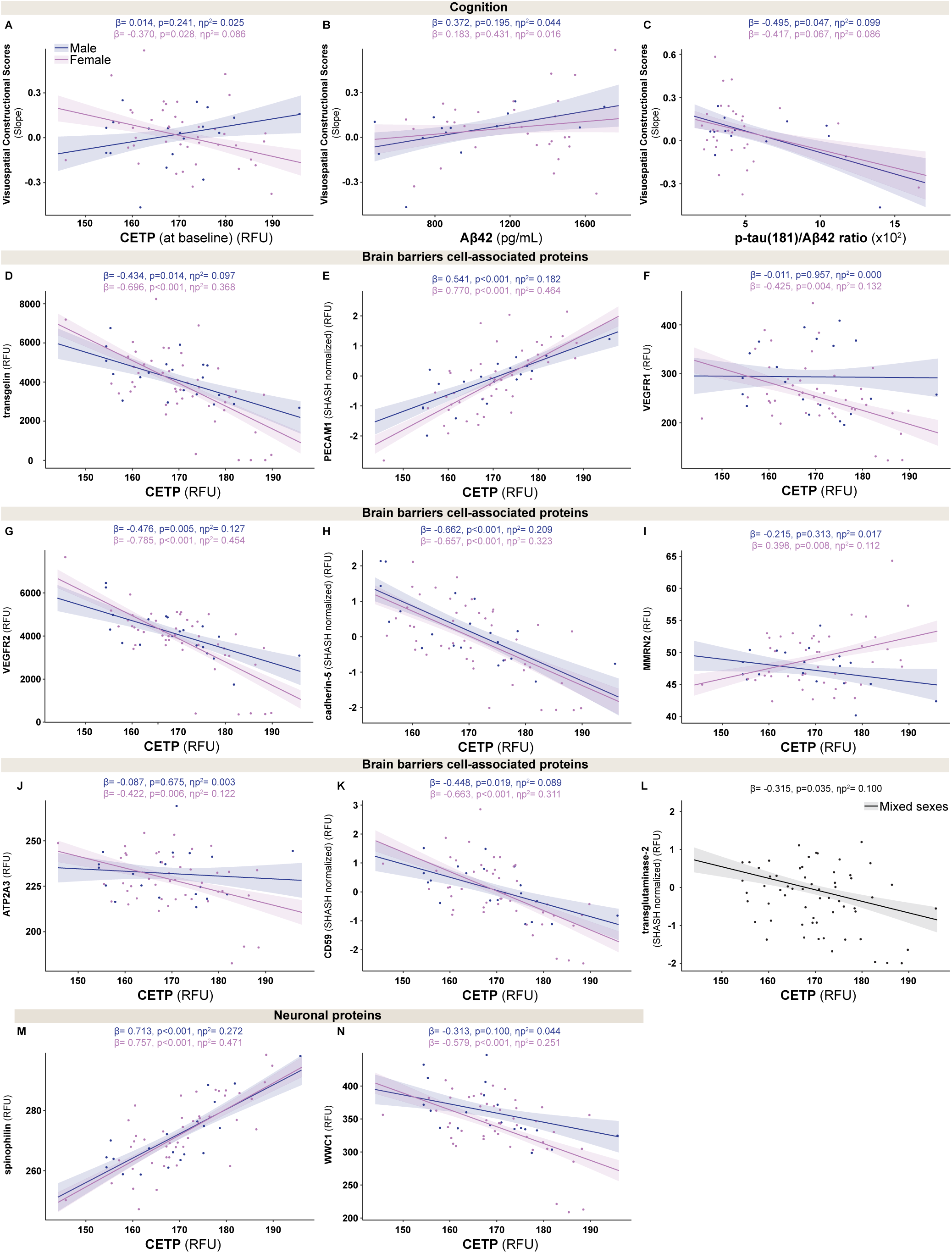
CSF analysis in human subjects corroborate findings from murine model. SOMAscan proteomics on human CSF of the PREVENT-AD cohort were performed and relationships were analyzed with cognitive scores and proteins of differentially regulated genes identified by RNAseq in the mouse model. **A-C** VCIS slopes (A), classic AD biomarker Aβ42 (B), and p-tau(181)/ Aβ42 ratio in relationship to baseline CETP levels in CSF. **D-N** CFS proteins From the SOMAscan data, brain barrier proteins including transgelin, VEGFR1, VEGFR2, cadherin-5, MMRN2, PECAM1, and CD59 were plotted over CSF CETP at baseline. **M, N** Additionally, the effect of baseline CSF CETP on neuronal protein spinophilin and WWC1 were analyzed. Data is shown as scatter plots with a linear regression line for the main effect of interest. Relationships were analyzed using GLMs and stratified by sex, controlling for participant age, *APOE* ε4 status, and years of education (cognition analyses only). Standard error of the regression line is shown on the plot as full lines surrounding the linear regression. **A-J, L, M** Data points from female subjects are represented in violet and from male subjects in indigo, with the regression line with the corresponding shade, *n=19-46*. **L** Data points from male and female subjects are represented in black with the regression line with the corresponding shade and were combined showing significance when pooled but not as separate analyses, *n=20-46*. **A-N** β-coefficient (β), p-value (p) and partial eta squared (ηp^2^) are denoted for each analysis.

## Discussion

Conventionally studied in cardiovascular research, CETP has now gained attention in neurodegenerative diseases. Several epidemiological studies have shown that low CETP activity reduces the risk of AD *^15, 20–27^*. Our research explored the effects of pharmacological CETP inhibition in an AD mouse model and compared these findings to the at-risk asymptomatic PREVENT-AD human cohort. Our study aimed to determine whether cognitive decline could be prevented in an AD mouse model mitigated by CETP inhibition. Focusing on early disease development, we assessed the impact of CETP on cognition, AD biomarkers, as well as cholesterol regulation. Our findings showed that evacetrapib preserved memory in female mice independent of Aβ accumulation, aligning with human data*^86–89^*. Mechanistically, we demonstrate that evacetrapib preserves memory by altering plasma lipoproteins, supporting cerebrovascular homeostasis, and optimizing brain lipid distribution. Whether CETP inhibition in the periphery, the brain, or both contributed to memory preservation remains to be determined*^37, 45, 90^*. The results of our study strongly indicate that inhibiting CETP activity during the early stages of AD has significant potential to prevent or delay memory decline*^18^*.

Remarkable are the sex differences in our mouse model and in part in the human data. Our results showed significant memory maintenance and modulation of plasma lipoproteins in female mice, with attenuated effects in males. Several other mouse studies investigating lipid changes between sexes reported as well that alterations in female mice lead to more profound responses to lipoprotein changes compared to their male counterparts in line with our observations*^91, 92^*. We conclude that that the attenuated effects in evacetrapib-treated male mice is likely due to their diminished modulation of the lipoprotein profile rather than to a limitation of our model. Additionally, sex differences in AD are well-established, with women being twice as likely to develop the condition*^93^*. Most if not all women affected by AD are post-menopausal, and are particularly at risk, often exhibiting lower HDL and higher LDL levels compared to men*^36, 94–96^*. This disparity in risk can be attributed to the influence of estrogen: elevated estrogen levels enhance HDL, and both estrogen and HDL play protective roles in the vasculature, which diminish with the onset of menopause*^97–99^*. Collectively, these findings highlight that lipoprotein changes in females, which can be modulated by CETP inhibitors, have a strong impact on health and cognition, especially after menopause in elderly women.

A striking observation in our study is that the effects of CETP on cognition are independent of Aβ indicating that other factors contribute to the causes of memory impairment in AD. Similarly, Tong et al. demonstrated in a preclinical study investigating statins, a drug class inhibiting cholesterol synthesis, that lipid modulation improved cognitive function in 6-month-old APPJ20 mice independent of Aβ species or plaque load*^100^*. Notably, the demonstrated efficacy of lipid-modifying agents on cognition has primarily been shown in younger subjects*^101^*, a phenomenon also observed in mice*^100, 102^*. Data from the PREVENT-AD human cohort investigating the early progression into dementia support these findings, showing no relationship between conventional AD markers Aβ42 or the p-tau181/Aβ42 ratio with CSF CETP abundance or cognition in women replicating the findings in our mouse model. Furthermore, in asymptomatic PREVENT-AD subjects, decreased levels of Aβ42 in the CSF did not yet demonstrate a relationship with poorer cognition in either VCIS or overall RBANS slopes, which was previously shown before in other studies performed in symptomatic MCI or AD patients *^86–89^*. While reduced CSF Aβ42 is recognized as a marker of AD, our study focuses on the subjects/patients at their pre-clinical pre-MCI stage*^103^*. Importantly, our results, consistent with our young mouse model, showing that cognitive slopes over time, more specifically the prediction of cognitive impairment, was associated with CETP abundance in the CSF at baseline in female subjects before disease onset corroborating the importance of CETP in AD and relevance of our mouse model to human physiology.

The involvement of lipid metabolism and the potential of pharmacological lipid interventions as disease modulators in dementia is not new*^104–107^*. A meta-analysis investigating the impact of statin treatment, concluded a potential favorable role of statins on AD and dementia*^105^*. While statins may offer overall benefits for AD, their effectiveness is not universal. Women who carry the APOE4 genotype, which is prevalent among AD patients, are particularly impacted by the limitations of statins, as they typically exhibit a less favorable response to treatment compared to men*^108, 109^*. Furthermore, side effects associated with statin use have previously been noted, such as memory loss and confusion*^110^*. Mechanistically, these adverse effects may stem from the inhibition of HMGCR, the rate-limiting enzyme in the mevalonate pathway*^111–113^*. Although this pathway is primarily responsible for sterol synthesis, it also produces isoprenyl intermediates, which serve as critical posttranslational modifications for the function of small GTPases*^113^*. Emerging evidence suggests that prenylated proteins, particularly small GTPases, play significant roles in cognition and AD pathogenesis*^65, 114^*. Consequently, while statins have a positive impact on AD, they are not the ideal choice for standard treatment, especially once diagnosis is formally established. In this context, pharmacological CETP inhibition is a promising alternative, particularly in the context of presymptomatic prevention. New-generation CETP inhibitors like evacetrapib and obicetrapib have been proven to be safe, well-tolerated, and effective in lowering CETP activity in cardiovascular clinical trials*^10, 11, 13, 14^*. Epidemiological studies have highlighted the benefits of low CETP activity, especially at early AD stages in APOE4 carriers providing a significant advantage over statins*^21–27^*. Given that CETP is a highly specific enzyme, adverse effects associated with statins are unlikely to occur with CETP inhibition*^1, 115, 116^*. Additionally, CETP inhibitors offer potential benefits for various other diseases. They reduce the risk of type 2 diabetes by increasing HDL-C, which is inversely related to diabetes risk*^16, 117^*, and notably type 2 diabetes increases the risk for AD*^118^*. Ultimately, as outlines in the introduction, decreased CETP activity is linked to various health benefits such as protection from PD, ALS, sepsis, and cancer along with AD*^15^*. Thus, CETP inhibitors as a preventive treatment hold significant promise for AD and may additionally offer protective benefits against several other conditions, enhancing their potential as a novel therapeutic approach.

We observed significant but modest transcriptional changes between evacetrapib- and vehicle-treated CETP-expressing mice (24 genes), which could be attributable to using bulk mRNA extracted from total brain tissue. The majority of DEGs between evacetrapib and vehicle-treated CETP-expressing mice are expressed by endothelial cells or VSMCs and are involved in angiogenesis and vascular homeostasis*^71–73^*. Notably, single-cell RNA sequencing of human brain has revealed that CETP is almost exclusively expressed in brain endothelial cells*^2^*. Brain endothelial cells only constitute 0.2-7% of all brain cells, thus the observed changes indicate that evacetrapib has profound impacts on the brain vasculature*^119^*. Previous mouse studies have linked *CETP* expression to endothelial dysfunction through nitric oxide synthase (NOS) mechanisms, suggesting that CETP inhibition by evacetrapib maintains cerebrovascular integrity*^120, 121^*. In the PREVENT-AD cohort, our murine results align with the human data where several endothelial-specific and -related proteins were also identified in the CSF of subjects showing significant relationships with CETP levels. While the proteins encoded by these genes promote vascular stability and angiogenesis in tissues, the relationship between their levels in CSF and brain tissue remains to be investigated*^74, 78, 122–126^*. Nevertheless, CSF changes for several proteins have been characterized*^127–130^*. For instance, disruptions in the choroid plexus epithelium and enzymes like Na-K-ATPase, of which ATP2A3 is a part of, have been observed in the CSF of AD patients*^128^*. Additionally, decreased CSF levels of transgelin-2, a protein encoded by a gene paralog to *TAGLN*, and VEGFR2 have been associated with conditions such as AD and Amyotrophic Lateral Sclerosis (ALS), while elevated CSF PECAM-1 levels have been reported in multiple sclerosis, emphasizing the importance of endothelial proteins changes in neurodegeneration and early AD*^127, 129, 130^*. The overlap between regulated genes and their protein products suggests that similar mechanisms might be at play in both our mouse model and at-risk asymptomatic humans, emphasizing the role of CETP in cognitive function, endothelial health and brain barrier functions independent of traditional AD biomarkers.

Neuronal loss in the hippocampus is a well-documented process in AD*^131^*. We demonstrated that evacetrapib increased free cholesterol in the dentate gyrus region of the hippocampus. Concurrently, our RNA sequencing revealed alterations in neuronal genes, suggesting that redistribution of lipids in hippocampus may drive these neuronal changes, with cholesterol as a key factor. Cholesterol is critical for synaptogenesis, forming lipid rafts that cluster synaptic proteins and stabilize synapses, indicating a potential link between lipid distribution and neuronal function in the brain*^132^*. Several studies have previously highlighted that a reduction in free cholesterol in the hippocampus impairs synaptic proteins mobility, emphasizing its crucial role in preserving synaptic health and cognitive function*^132–135^*. Thus, the redistribution of lipids, particularly cholesterol, in the hippocampus may contribute to neuronal changes, emphasizing its essential role in maintaining synaptic integrity and good cognition.

In summary, these findings position CETP as a highly promising drug target in AD, with significant potential for repurposing CETP inhibitors like evacetrapib. Our research demonstrates that increased CETP expression impairs cognition, while pharmacological inhibition of CETP preserved cognitive function. The beneficial effects of CETP inhibition appear to be two-fold: in the brain, it increases cholesterol levels in the hippocampus potentially maintaining neuronal function; in the periphery, it modifies plasma lipoproteins thereby improving cerebrovascular health. Importantly, we established that our findings in our mouse model are highly relevant to human physiology, enhancing the scientific validity of our work. We assert that prophylactically administered CETP inhibitors such as evacetrapib before disease onset could effectively preserve memory and delay or prevent the progression of AD.

## Author contributions

JP contributed to the experimental design, breeding and mouse handling, conducting experiments, and interpretation of results, and draft of the original manuscript. IS and JP^a^ contributed to the analyses for the human subjects data, interpretation and contributed to the original draft of the manuscript. MSK and JP contributed to the preparation of libraries. MSK, HN, and WAP contributed to the RNA sequencing analysis and interpretation libraries and original draft of the manuscript. RSK contributed to the interpretation of results. AK and DL designed and performed the GC-MS analysis of lipids. LMM, JP^a^, and WAP, designed and supervised the project.

## Acknowledgements

We thank the donors and their families for the use of human CSF tissue in this study and staff of the Douglas Research Center, Montreal, Canada, and the PREVENT-AD for making it available. We would like to express our sincere gratitude to Dr. Claudio Cuello, McGill University, for generously providing the McGill-APP-Thy1 mice. Images were aquired and analyzed using the McGill University Advanced BioImaging Facility (ABIF) (RRID:SCR_017697). The Comparative Medicine and Animal Resources Centre (CMARC) and Dr. Paméla Thebault from the Immuno-monitoring Clinic Platform at the Centre Hospitalier de l’Université de Montréal (CHUM), and Nailya Ismailova and Dr. Jorg Fritz from McGill University, all contributed to the work. We thank Anne Labonté and Laurence Maligne-Bruneau for their technical contribution in the preparation and analyses of the human CSF samples from the PREVENT-AD cohort.

## Consent Statement

Each PREVENT-AD participant and their study partner provided written informed consent. All procedures were approved by the McGill University Faculty of Medicine Institutional Review Board and complied with the ethical principles of the Declaration of Helsinki.

## Supplemental figure legends

**Supplemental Figure 1.**
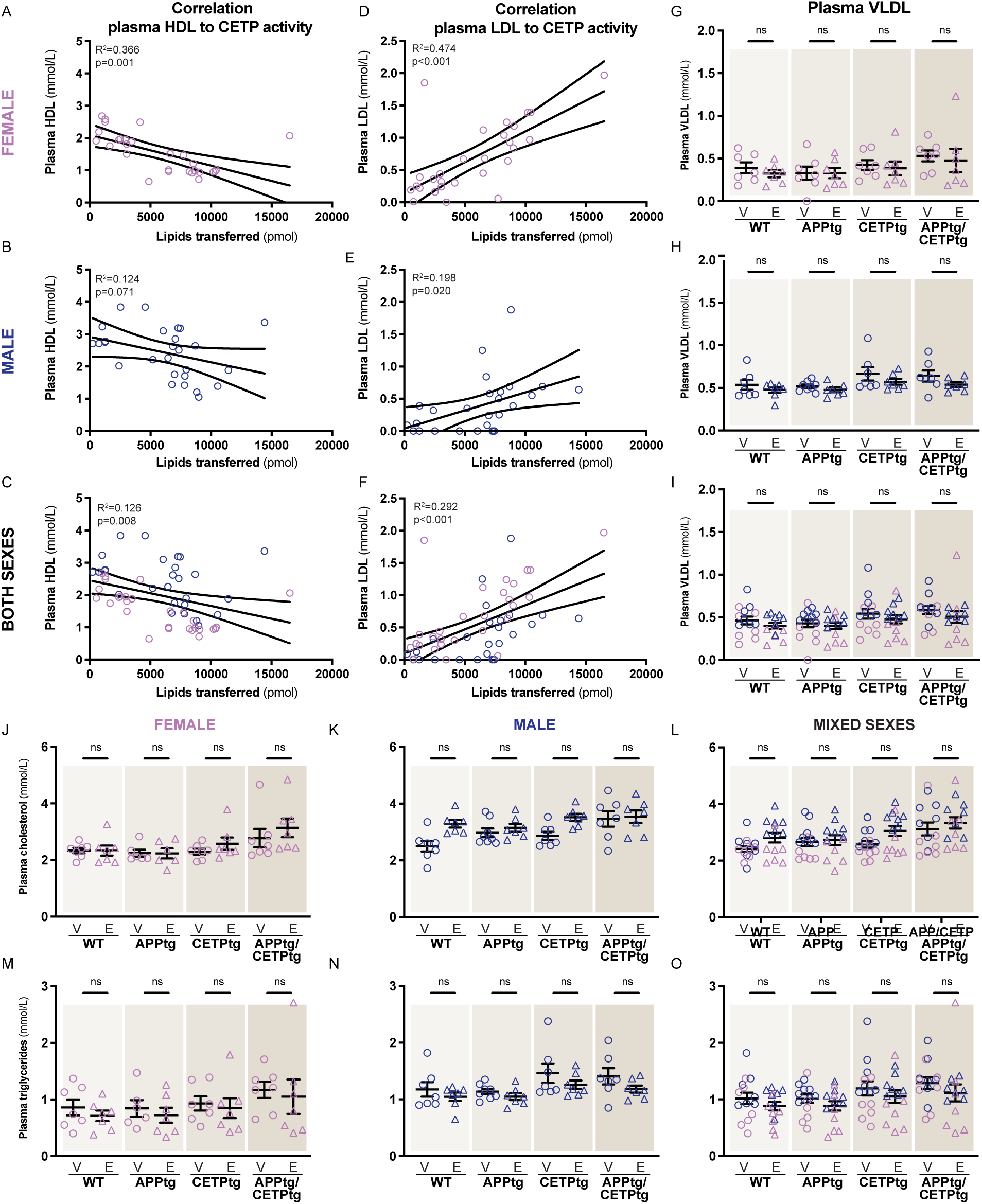
Correlations with plasma lipoproteins and CETP activity, VLDL levels, as well as total plasma cholesterol and triglycerides. **A-F** Linear correlations were calculated to compare CETP activity plotted against HDL and LDL levels, *n*=*27-28*, levels in female, male and mixed sexes cohorts of mice. Data is shown as scatter plots with a simple linear regression line. Correlations were analyzed using Pearson’s correlation assuming normality and the 95% confidence interval is shown on the plot as dotted lines. R-squared (R^2^) and p-values (p) are indicated for each correlation. **G-I** Plasma lipoprotein VLDL, **J-L** total cholesterol, and **M-O** total triglycerides levels were quantified in female and male mice, *n*=*6-8,* as well as both sexes combined, *n=12-14*. **G-O** Data is shown as bar charts with individual points. Circles represent the vehicle condition while triangles denote the evacetrapib condition. Bar symbols indicate mean ± SEM, where female data points are represented in violet and male data points in indigo. Statistical significance was measured using two-way ANOVA followed by Tukey’s post-hoc test.

**Supplemental Figure 2.**
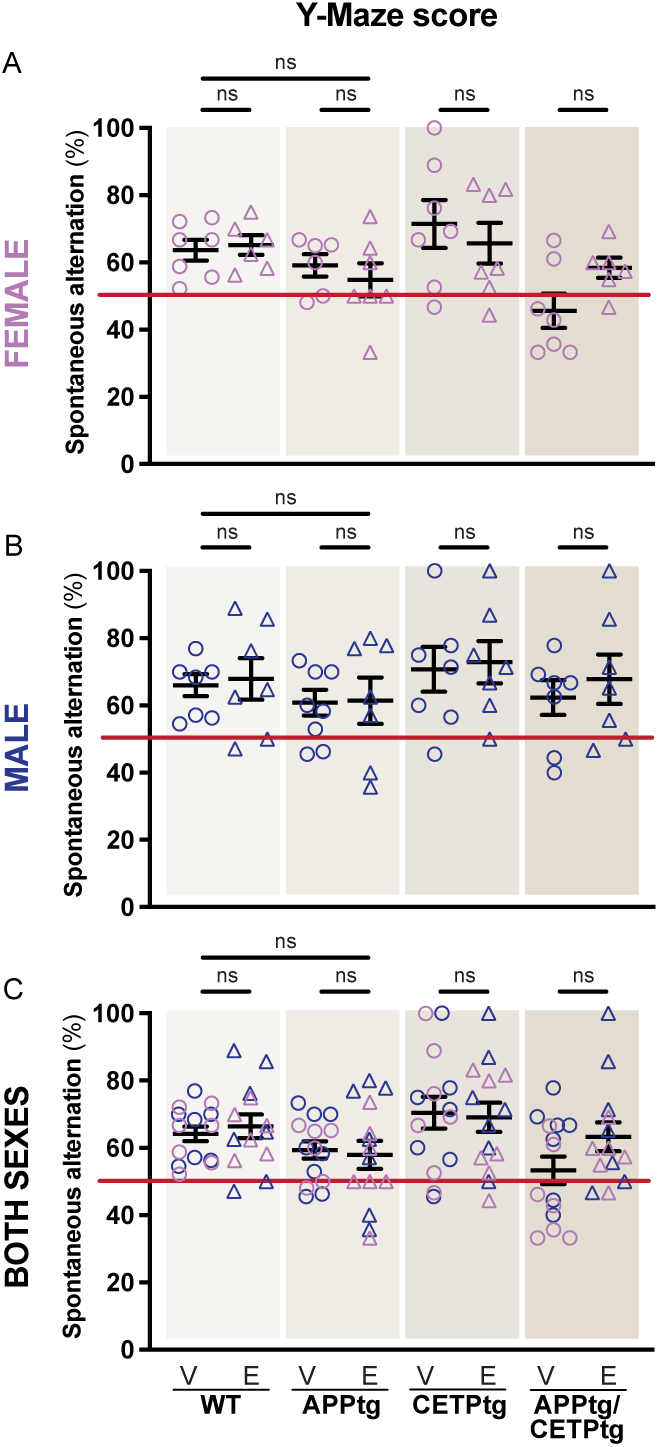
Y-maze test. **A-C** Y-maze scores were displayed for female and male mice, *n*=*6-8,* as well as both sexes combined, *n=12-14*, with a red line indicating the 50% discrimination ratio at which mice cannot differentiate between the familiar and the novel object suggesting memory impairment. Circles represent the vehicle condition while triangles denote the evacetrapib condition. Bar symbols indicate mean ± SEM, where female data points are represented in violet and male data points in indigo. Statistical significance was measured using two-way ANOVA followed by Tukey’s post-hoc test.

**Supplemental Figure 3.**
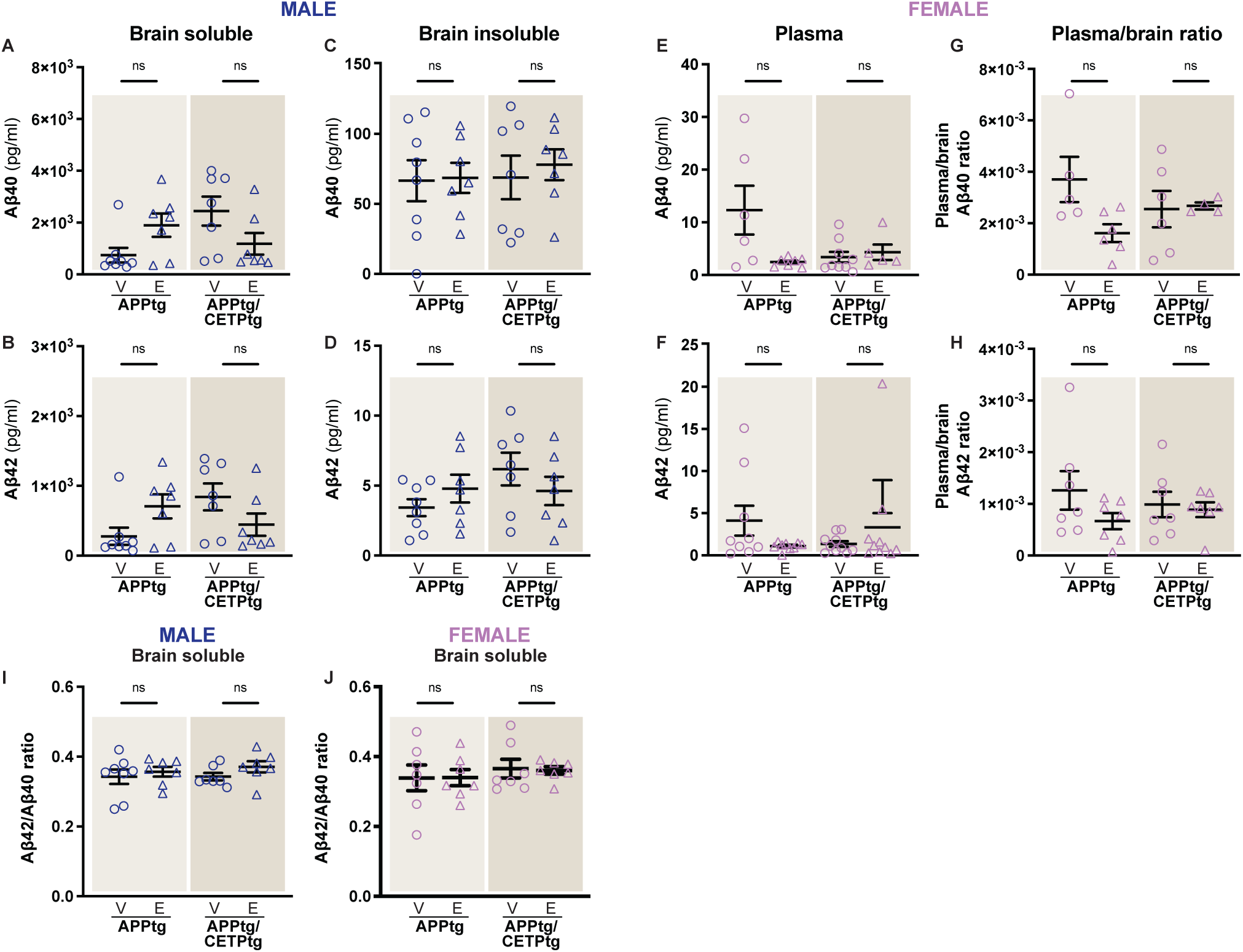
Aβ40 and Aβ42 quantification in male brains, female plasma, as well as plasma to brain and Aβ42/Aβ40 ratios. **A-D** Levels of soluble and insoluble Aβ40 and Aβ42 peptides were quantified in brain homogenates of male mice using ELISA. **E, F** Aβ40 and Aβ42 peptides were quantified in plasma of female mice using ELISA. **G, H** Plasma to brain ratio for Aβ40 and Aβ42 were plotted to identify potential blockages in excretion. **I, J** Aβ40/Aβ42 ratio plotted in male and female for soluble Aβ peptides. **A-J** Data is shown as bar charts with individual points. Circles represent the vehicle condition while triangles denote the evacetrapib condition. Bar symbols indicate mean ± SEM, *n*=*6-8,* where female data points are represented in violet and male data points in indigo. Statistical significance was measured using two-way ANOVA followed by Tukey’s post-hoc test.

**Supplemental Figure 4.**
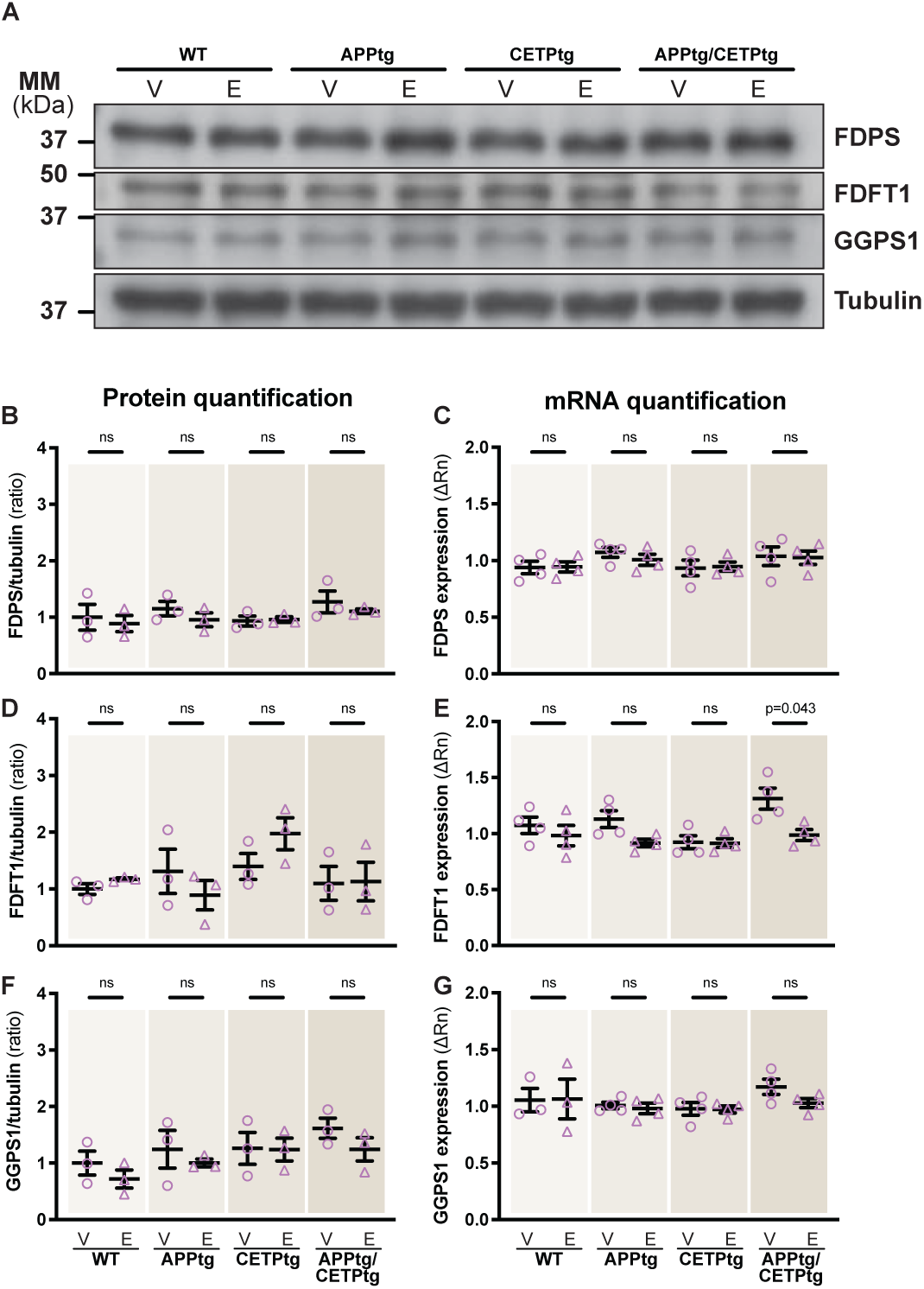
Enzymes in the mevalonate/isoprenoid pathway in brain tissue. **A** Female mouse brain was compared for enzymes FDPS, FDFT1, GGPS1, and tubulin as loading control. Representative images of three biological replicates are shown for each immunoblot panels. Protein quantification from western blot and mRNA quantification was done for **B, C** FDPS, **D, E** FDFT1, and **F, G** GGPS1. Data is shown as bar charts with individual points. Circles represent the vehicle condition while triangles denote the evacetrapib condition. Bar symbols indicate mean ± SEM, *n=3-4*, where female data points are represented in violet. Statistical significance was measured using two-way ANOVA followed by Tukey’s post-hoc test.

**Supplemental Figure 5.**
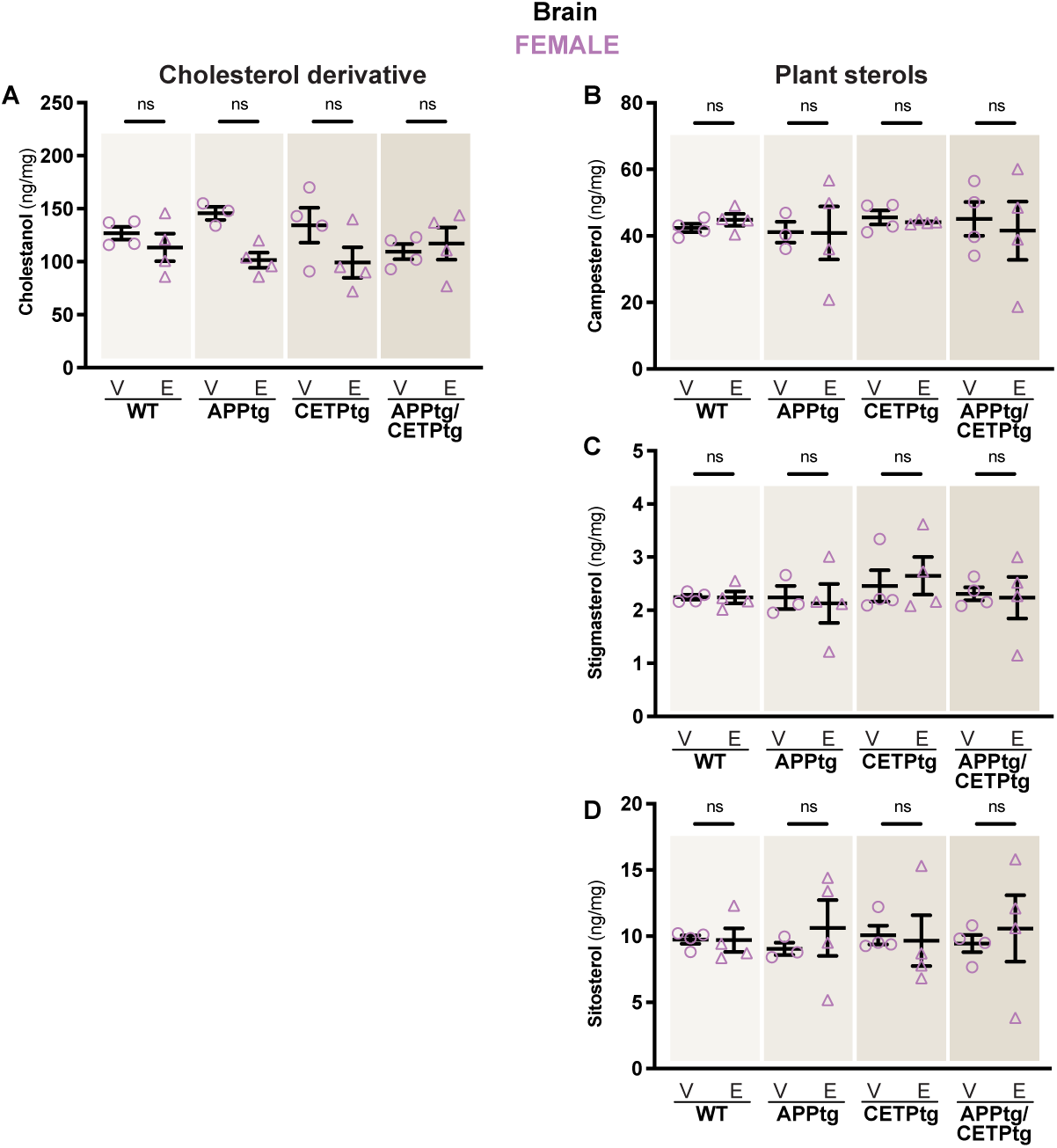
Food-derived plant sterols and cholesterol derivative in brain tissue. **A** Cholestanol and **B-D** Plant sterols campesterol, stigmasterol and sitosterol were quantified using GC-MS from brain tissue from female mice. **A-D** Data is shown as bar charts with individual points. Circles represent the vehicle condition while triangles denote the evacetrapib condition. Bar symbols indicate mean ± SEM, *n=3-4*, where female data points are represented in violet. Statistical significance was measured using two-way ANOVA followed by Tukey’s post-hoc test.

**Supplemental Figure 6.**
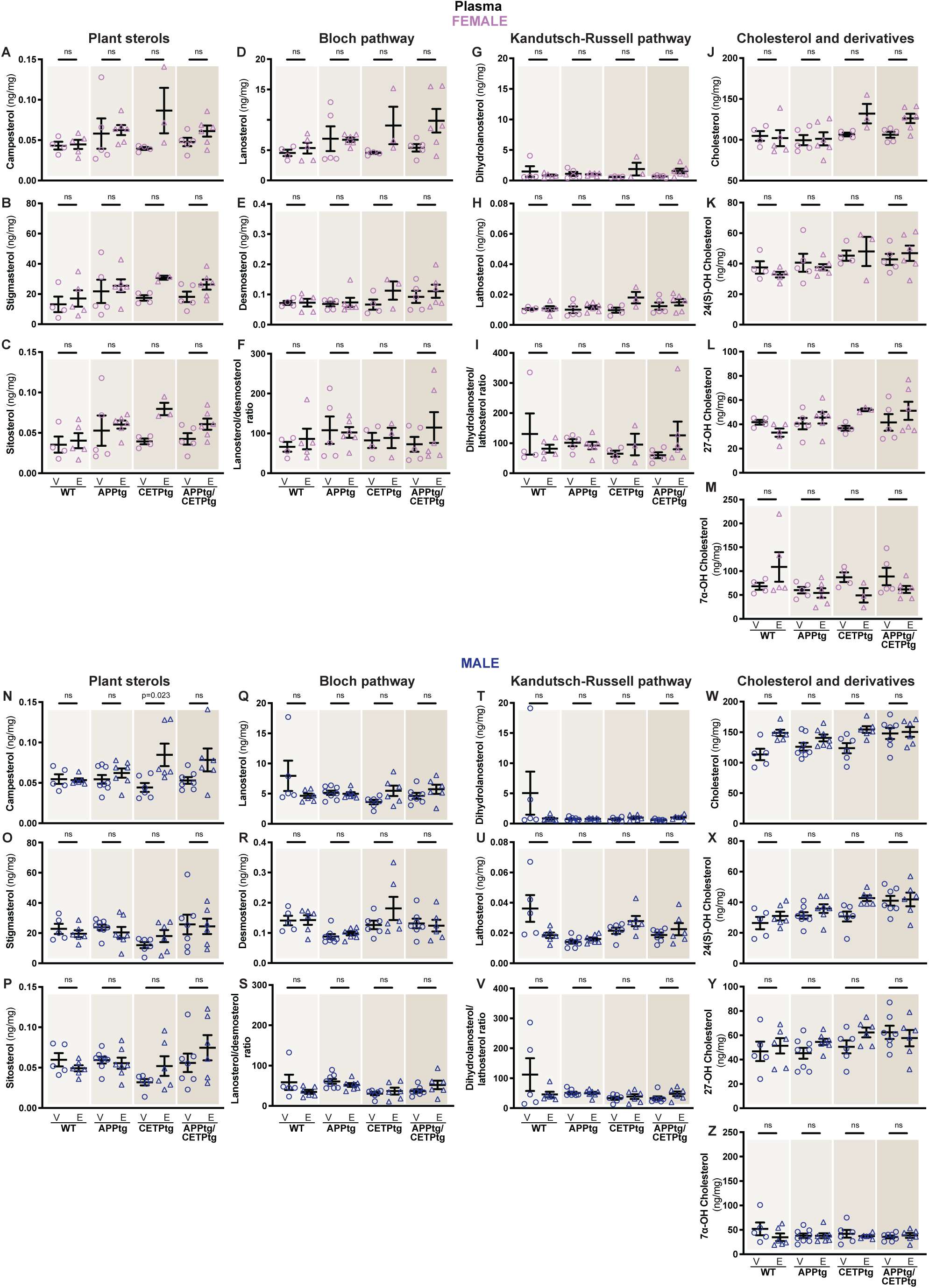
Plant sterols, cholesterol precursors, cholesterol, and cholesterol derivatives analysis in plasma. Cholesterol analyses from plasma of **A-M** female and **N-Z** male mice. Analysis includes plant sterols campesterol, stigmasterol and sitosterol. cholesterol precursors lanosterol and desmosterol, dehdrolanosterol and lathosterol, total cholesterol and cholesterol derivatives 24(S)-hydroxycholesterol (24(S)-OH), 27-hydroxycholesterol (27-OH), and 7α-hydroxycholesterol (7α-OH). **F, I, S, V** Ratios of lanosterol to desmosterol, and 24.25-dihydrolanosterol to lathosterol were plotted. **A-Z** Data is shown as bar charts with individual points. Circles represent the vehicle condition while triangles denote the evacetrapib condition. Bar symbols indicate mean ± SEM, *n=6-8*, where female data points are represented in violet and male data points in indigo. Statistical significance was measured using two-way ANOVA followed by Tukey’s post-hoc test.

**Supplemental Figure 7.**
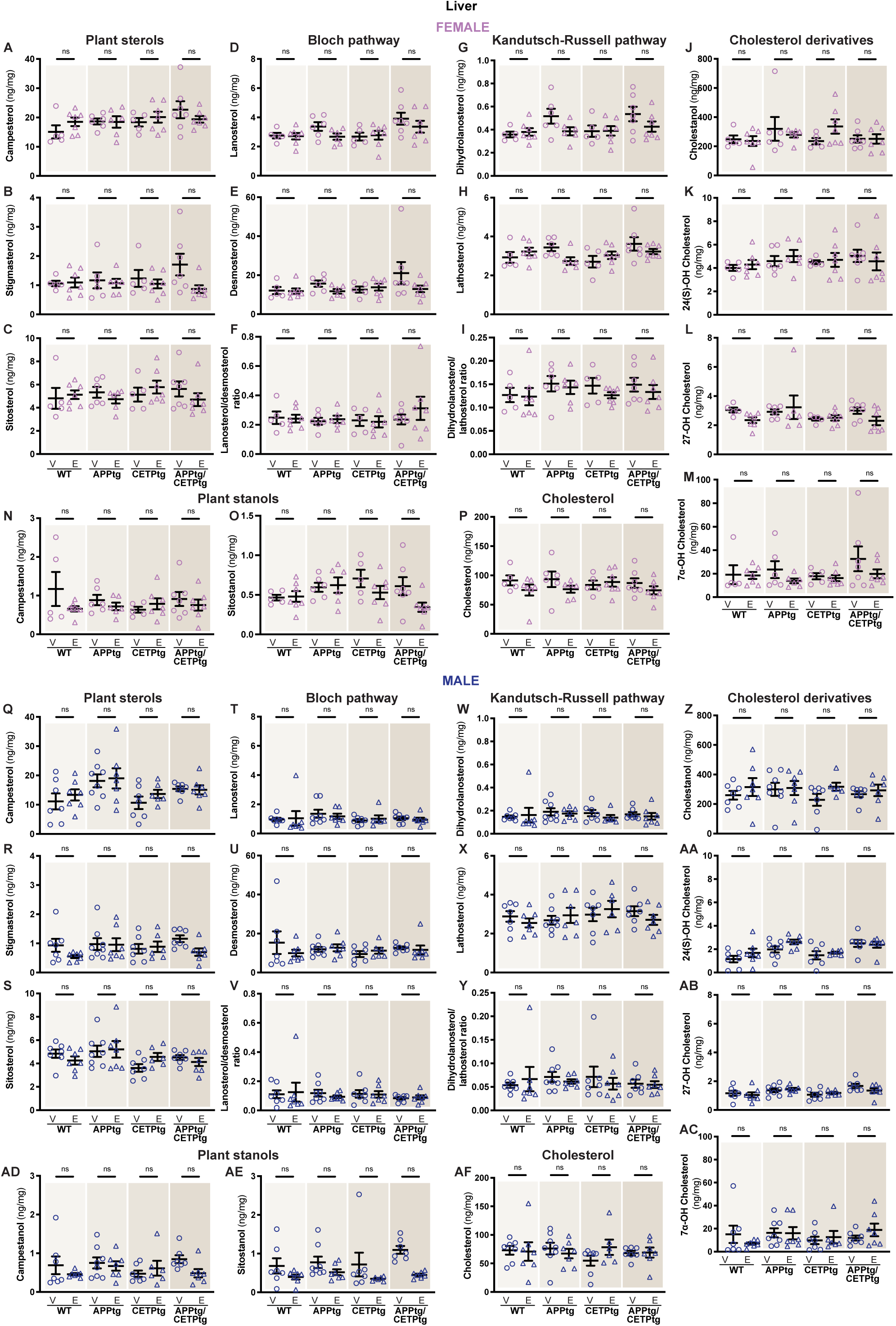
Plant sterols, plant stanols, cholesterol precursors, cholesterol, and cholesterol derivatives analysis in liver tissue. **A-AF** Cholesterol analyses from liver tissue of female (A-P) and male (Q-AF) mice. Analysis includes plant sterols campesterol, stigmasterol and sitosterol. cholesterol precursors lanosterol and desmosterol, dehdrolanosterol and lathosterol, total cholesterol and cholesterol derivatives 24(S)-hydroxycholesterol (24(S)-OH), 27-hydroxycholesterol (27-OH), and 7α-hydroxycholesterol (7α-OH). **F, I, V, Y** Ratios of lanosterol to desmosterol, and 24.25-dihydrolanosterol to lathosterol were plotted to identify potential blockages in the pathway. **A-AF** Data is shown as bar charts with individual points. Circles represent the vehicle condition while triangles denote the evacetrapib condition. Bar symbols indicate mean ± SEM, *n=6-8*, where female data points are represented in violet and male data points in indigo. Statistical significance was measured using two-way ANOVA followed by Tukey’s post-hoc test.

**Supplemental Figure 8.**
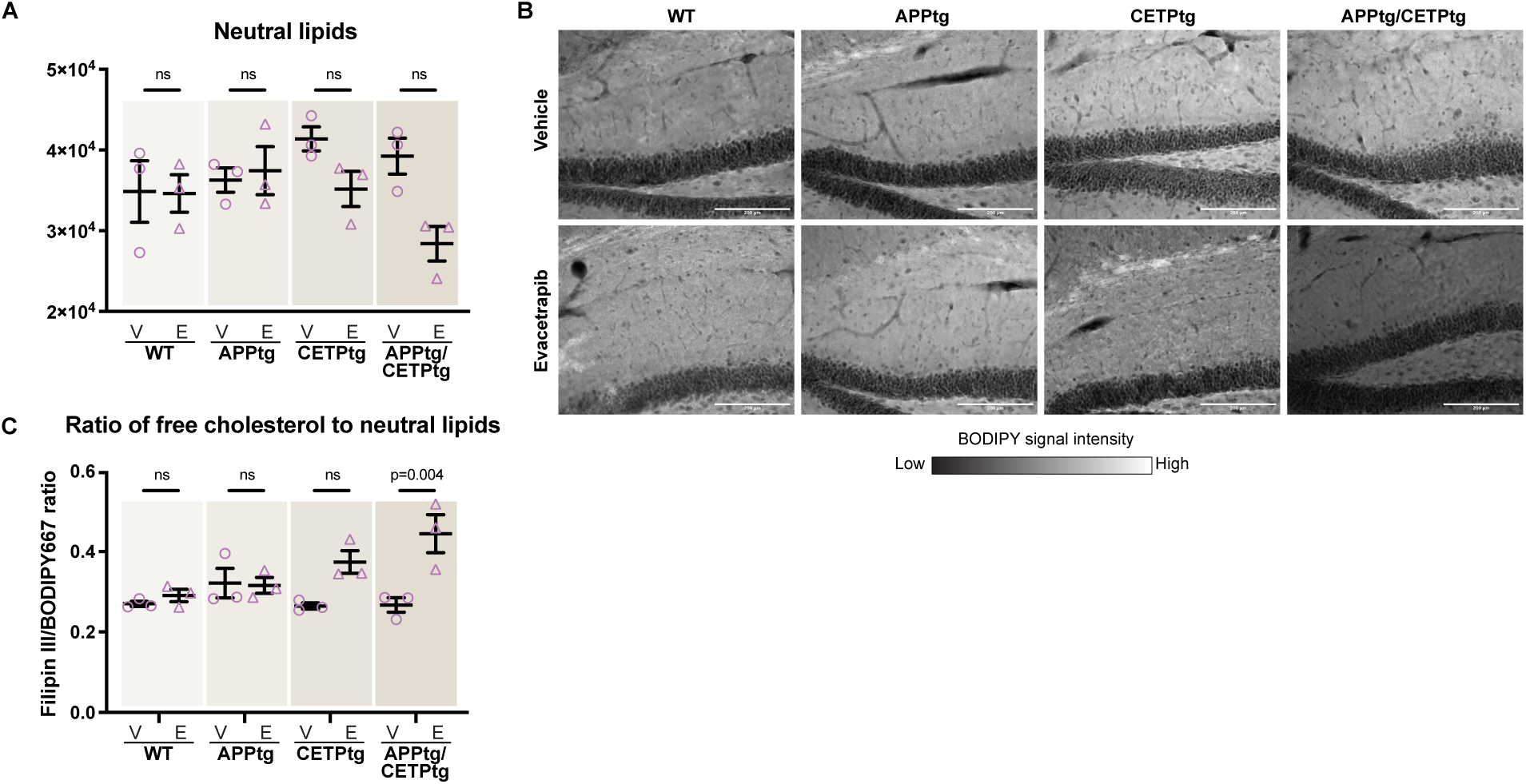
Neutral lipids distribution in the dentate gyrus region of the hippocampus. **A, B** Quantification of BODIPY665/676 staining in the dentage dyrus of the hippocampus of female mice. **B** Representative images of sagital sections at 20X magnification in female mouse brains with signal intensity is depicted using a scale bar, with lighter shades representing higher intensity and darker shades indicating lower intensity. Representative images of three biological replicates are shown for each microscopy panel. **C** Free cholesterol to neutral lipids signal is represented as a ratio. **A, C** Data is shown as bar charts with individual points. Circles represent the vehicle condition while triangles denote the evacetrapib condition. Bar symbols indicate mean ± SEM, *n=3*, where female data points are represented in violet. Statistical significance for all analyses was measured using two-way ANOVA followed by Tukey’s post-hoc test.

**Supplemental Figure 9.**
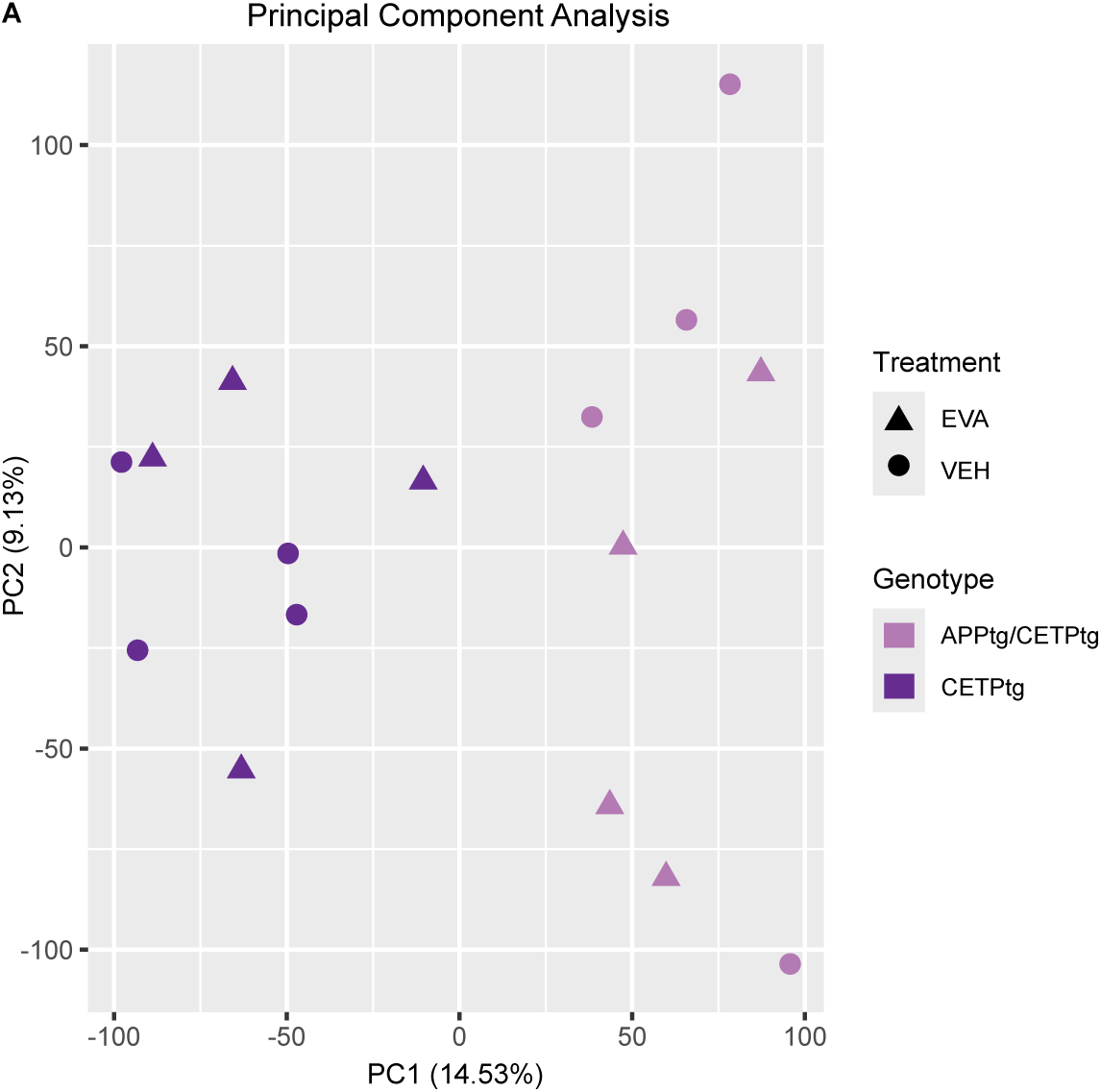
Principal component analysis. **A** PCA analysis. Principal components PC1 and PC2 are shown with 14.53% and 9.13% variance explained, respectively. Circles represent the vehicle condition while triangles denote the evacetrapib condition. Dark purple is set for CETPtg and violet for APPtg/CETPtg replicates.

**Supplemental Figure 10.**
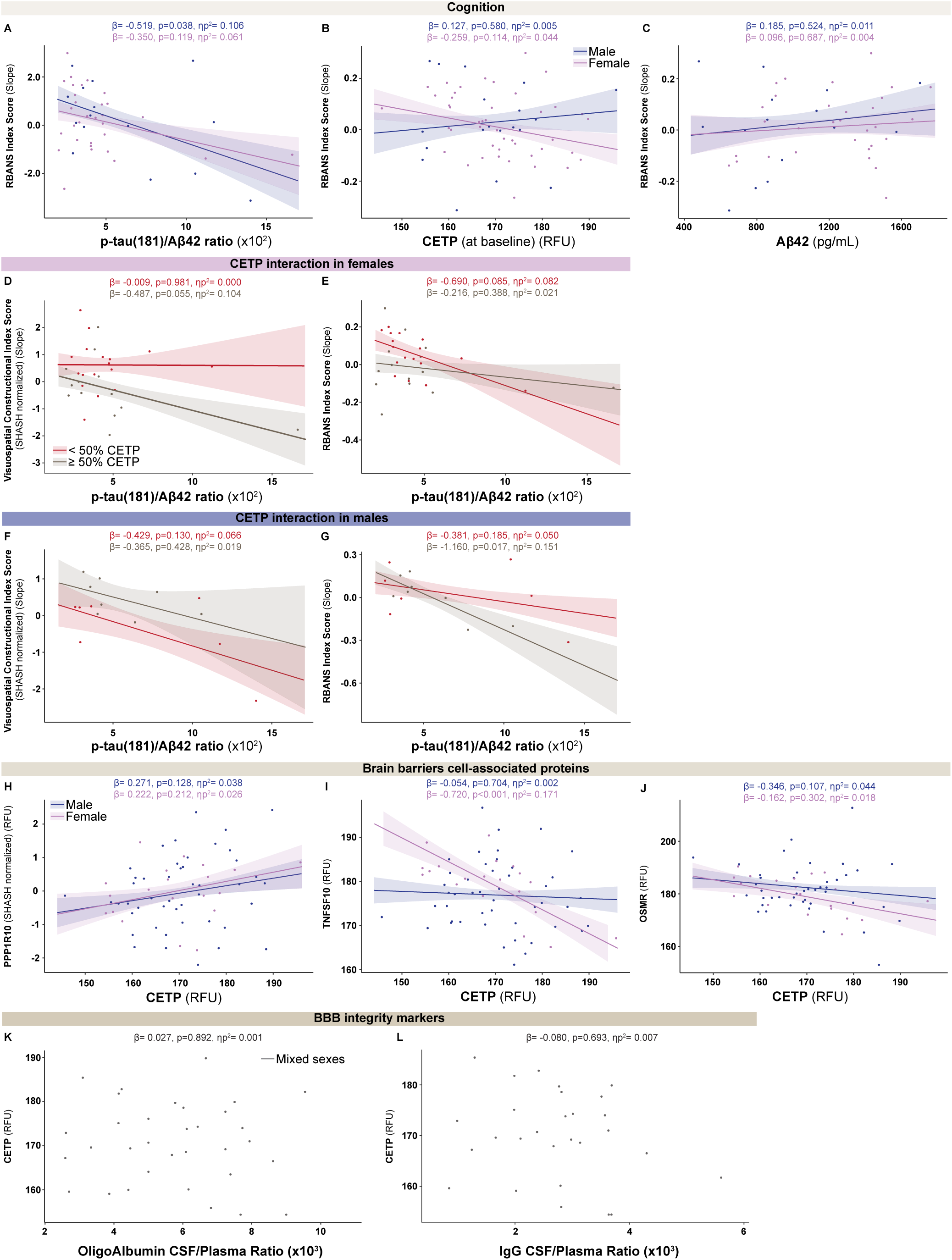
CSF CETP effects on cognition and CSF proteins. **A-L** SOMAscan analysis of CSF from human subjects *n=7-46*. **A-C** RBANS slopes in relationship to p-tau(181)/Aβ42 ratio, CETP levels in CSF, and classic AD biomarker Aβ42 were plotted. **D-G** Relationship between three parameters, p-tau(181)/Aβ42, RBANS or VCIS, and CETP showing either < 50% and ≥ 50% top and bottom CETP expressors. **H-J** From the SOMAscan data, brain barrier proteins and neuronal proteins including PPP1R10, TNFSF10, and OSMR in relation to CSF CETP. **K, L** Plasma to brain albumin and IgG ratio were analyzed to investigate BBB integrity. Data is shown as scatter plots with a simple linear regression line. Relationships were analyzed using GLM stratified by sex, controlling for participant age, *APOE* ε4 status, and years of education (cognition analyses only). Standard error of the regression line is shown on the plot as full lines surrounding the linear regression. **A-L** Data points from female subjects are represented in violet and from male subjects in indigo, with the regression line with the corresponding shade. β-coefficient (β), p-value (p) and partial eta squared (ηp^2^) are denoted for each relationship.

## Supplemental methods

### Y-maze test

An adapted version of the Y-maze test based on the method by Kraeuter et al. was performed in the mice at 21 weeks of age to assess short term spatial memory*^59^*. Mice were habituated to handling 3-days prior to the experiment. The experiment was carried out in a sound-proof dark room in an open field arena and with an infra-red video camera recording mouse behaviour. At the start of the experiment, mice was placed in the middle of the 3-arm maze. Mice were allowed to freely explore the maze for five minutes while being recorded and then placed back to their home cage. The Y-maze and the examiner PPE were cleaned with 70% ethanol thoroughly and dried before each test to avoid olfactory bias.

Spontaneous alteration was measured as follows:

### Spontaneous alteration (%) = Number of entries into a novel arm/ (total alterations – 2) * 100

Each arm was denoted by a number (1, 2, or 3). The series of numbers corresponding to each arm entry were noted for each mouse. The first entry was discarded to avoid directional bias by the examiner into the maze. An entry into a novel arm was defined as entering an arm (e.g., arm 1) after having previously entered two different arms (e.g., arm 2 and arm 3) in any order. Mice with less than 8 entries into any arms of the maze were excluded from the analysis. The data was analyzed and plotted on GraphPad Prism (GraphPad Software Inc., USA).

### BODIPY665/676 staining

For the BOPIDY665/676 staining, a similar protocol as for the filipin III was used. The difference is that the 25 μM-thick free-floating slices were incubated in 20 μM BODIPY665/676 in a 0.3% PBS-Triton X solution for 2 h. BODIPY665/676 was chosen over BODIPY493/503 to reduce the autofluorescence from brain slices in the green channel.

### Western blot

Sodium dodecyl sulfate buffer with dithioreithol at a final concentration 5% (v/v) was added to samples and boiled at (70°C, 5 min). Proteins were separated on 10% polyacrylamide gels with Precision Plus Protein Dual Color Standards (Bio-Rad, USA) used as reference. Proteins were transferred onto nitrocellulose Membranes, 0.45 µm, (Bio-Rad, USA) by tank blotting (Bio-Rad, USA) at (400 mA, 4°C 2 h). The primary antibodies, anti-FDPS dilution 1:1000 (Abcam, USA, Catalog #ab195046), anti-FDFT1 dilution 1:1000 (Abcam, USA, Catalog # ab189874), anti-GGPS1 dilution 1:1000 (Cell Signaling, Catalog # ab167168), and secondary antibodies dilutions 1:10,000 (horseradish peroxidase-conjugated anti-rabbit, Promega, Catalog # W4011), were used. Signals were recorded on the Amersham Imager 600 (GE Healthcare Life Sciences, USA).

### qPCR analysis

Reverse transcription from RNA to complementary deoxyribonucleic acid (cDNA) was performed using the Applied Biosystems High-Capacity cDNA Reverse transcription Kit following manufacturer instructions (Life Technologies, USA). Total mRNA from brain was determined with SYBR Green qPCR mastermix (BioRad, USA) quantification using mouse specific primers. GAPDH and RSP18 were utilised as internal controls. RIN factor was measured for samples ranging between 7.3 and 8.6 (Agilent Technologies, USA).

Primer sequences:

FDPS:

5’-GGTGGTTCAGTGTCTGCTACGA-3’, 3’-CGCCTCATACAGTGCTTTCACC-5’.

FDFT1:

5’-GGATGTGACCTCCAAACAGGAC-3’, 3’-CAGACCCATTGAGTTGGCACAC-5’.

GGPS1:

5’-CACAGGCATTTAATCACTGGCTG-3’, 3’-CGTCGGAGCTTTGAACTGTCTTC-5’.

GAPDH:

5’-CATCACTGCCACCCAGAAGACTG-3’, 3’ATGCCAGTGAGCTTCCCGTTCAG-5’.

RSP18:

5’-CGGAAAATAGCCTTCGCCATCAC-3’, 3’-ATCACTCGCTCCACCTCATCCT-5’

### Principal component analysis (PCA)

PCA was performed using plotPCA function of ggplot2 version 3.5.1 and rlog normalization of gene raw counts.

### Albumin and IgG CSF/Plasma Ratios as biomarkers of BBB integrity

All analyses were part of the clinical routine diagnostic assessment of the PREVENT-AD participants between February 2008 and September 2017. Albumin and IgG levels in CSF and plasma were analyzed using immunoturbidimetric assays (kits NK004.OPT and NK032.L.OPT) on an Optilite analyser (Thermofisher Scientific, USA) at the department of clinical biochemistry at the McGill University Health Centre.

## Appendix

**Supplemental Table 1:** Internal controls for cholesterol, sterol precursors, plant sterols, and oxysterols in 100 µL plasma or to a chloroform/methanol tissue extract (5 mL chloroform/methanol, 2:1, v/v, per 10 mg dry adipose tissue).

| **Reagent** | **Provider** | **Stock** |
| --- | --- | --- |
| 50 µg of 5α-cholestane | Sigma-Aldrich, USA | 50 µL of a 1 mg/ml stock solution in cyclohexane from Merck, USA |
| 1 µg of epicoprostanol | Sigma-Aldrich, USA | 10 µL of a 100 µg/mL stock solution in cyclohexane |
| 50 ng of racemic [23,23,24,25-2H<sub>4</sub>]24(R,S)-hydroxycholesterol | Medical Isotopes, USA |  |
| 100 ng 26.26.26.27.27.27-[2H<sub>6</sub>]-7α-hydroxycholesterol | Medical Isotopes, USA |  |
| 100 ng [16,16,17,20,22-2H<sub>5</sub>]- (25R)27-hydroxycholesterol | Sugaris, USA | 50 µL of a 2 µg/mL stock solution in toluene from Merck, USA |

**Supplemental Table 2:** TMSi derivatives of the non-cholesterol sterols and di-TMSi-derivatives of the oxysterols monitoring compounds and specific mass per change (m/z):

| Compound | m/z |
| --- | --- |
| Lathosterol | 458 (M<sup>+</sup>) |
| Desmosterol | 441 (M<sup>+</sup> -15, M<sup>+</sup> -CH<sub>3</sub>) |
| Lanosterol | 393 (M<sup>+</sup> -90-15, M<sup>+</sup> -OTMSi-CH<sub>3</sub>) |
| 24.25-dihydrolanosterol | 395 (M<sup>+</sup> -90-15, M<sup>+</sup> -OTMSi-CH<sub>3</sub>) |
| Campesterol | 472 (M<sup>+</sup>) |
| Sitosterol | 486 (M<sup>+</sup>) |
| Stigmasterol | 484 (M<sup>+</sup>) |
| Campestanol | 474 (M<sup>+</sup>) |
| Sitostanol | m/z 488 (M<sup>+</sup>) |
| 26.26.26.27.27.27-[<sup>2</sup>H<sub>6</sub>]-7α-hydroxycholesterol | 462 (M<sup>+</sup> -90, M<sup>+</sup> -OTMSi) |
| 7α-hydroxycholesterol | 456 (M<sup>+</sup> -90, M<sup>+</sup> -OTMSi) |
| [23,23,24,25-<sup>2</sup>H<sub>4</sub>]-24(R,S)-hydroxycholesterol | 416 (M<sup>+</sup> -90-44, M<sup>+</sup> -OTMSi-CD(CH<sub>3</sub>)<sub>2</sub>) |
| 24(S)-hydroxycholesterol | 413 (M<sup>+</sup> -90-43, M<sup>+</sup> -OTMSi-CH(CH<sub>3</sub>)<sub>2</sub>) |
| [16,16,17,20,22-<sup>2</sup>H<sub>5</sub>]-27-hydroxycholesterol | 461 (M<sup>+</sup> -90) |
| (25R)-27-hydroxycholesterol | 456 (M<sup>+</sup> -90) |

**Supplemental Table 3:** Figure 6 detailed statistical analyses. **N**= sample size **β**= regression coefficient **SE**= standard error **p**= p-value **β 95% CI**= 95% confidence interval for β coefficient **pη^2^** = effect size

| Figure 6 | Predictor/Interaction | N | β | SE | p | β 95% CI | pη<sup>2</sup> |
| --- | --- | --- | --- | --- | --- | --- | --- |
| **A**<br>**(VCIS)** | CETP | 61 | -0.153 | 0.003 | 0.275 | [-0.431, 0.125] | 0.022 |
|  | CETP*Sex | 61 | -0.640 | 0.005 | 0.026 | [-1.199, -0.080] | 0.089 |
| **B**<br>**(VCIS)** | Aβ42 | 45 | 0.262 | 0.000 | 0.149 | [-0.098, 0.623] | 0.053 |
|  | Aβ42*Sex | 45 | -0.186 | 0.000 | 0.615 | [-0.929, 0.557] | 0.007 |
| **C**<br>**(VCIS)** | p-tau(181)/Aβ42 ($\times 10^2$) | 45 | -0.452 | 0.010 | 0.012 | [-0.799, -0.105] | 0.151 |
| | p-tau(181)/Aβ42 ($\times 10^2$)<br>*Sex | 45 | 0.079 | 0.018 | 0.798 | [-0.539, 0.696] | 0.002 |
| **D**<br>**(TAGLN)** | CETP | 66 | -0.612 | 162.532 | <0.001 | [-0.807, -0.147] | 0.393 |
|  | CETP*Sex | 66 | -0.263 | 346.725 | 0.211 | [-0.678, 0.153] | 0.026 |
| **E**<br>**(PECAM1)** | CETP | 65 | 0.692 | 0.009 | <0.001 | [0.516, 0.868] | 0.508 |
|  | CETP*Sex | 65 | 0.229 | 0.019 | 0.218 | [-0.139, 0.596] | 0.026 |
| **F**<br>**(VEGFR1)** | CETP | 66 | -0.291 | 0.784 | 0.016 | [-0.527, -0.057] | 0.092 |
|  | CETP*Sex | 66 | -0.413 | 1.657 | 0.100 | [-0.909, 0.082] | 0.044 |
| **G**<br>**(VEGFR2)** | CETP | 66 | -0.686 | 12.727 | <0.001 | [-0.871, -0.500] | 0.473 |
|  | CETP*Sex | 66 | -0.308 | 26.960 | 0.121 | [-0.701, 0.084] | 0.040 |
| **H**<br>**(CDH5)** | CETP | 65 | -0.659 | 0.010 | <0.001 | [-0.856, -0.462] | 0.428 |
|  | CETP*Sex | 65 | 0.005 | 0.022 | 0.980 | [-0.413, 0.423] | 0.000 |
| **I**<br>**(MMRN2)** | CETP | 65 | 0.200 | 0.050 | 0.112 | [-0.048, 0.449] | 0.041 |
|  | CETP*Sex | 65 | 0.612 | 0.104 | 0.020 | [0.099, 1.126] | 0.088 |
| **J**<br>**(ATP2A3)** | CETP | 64 | -0.308 | 0.186 | 0.014 | [-0.551, -0.065] | 0.098 |
|  | CETP*Sex | 64 | -0.335 | 0.387 | 0.190 | [-0.842, 0.171] | 0.029 |
| **K**<br>**(CD59)**<br>**(Robust errors)** | CETP | 66 | -0.594 | 0.011 | <0.001 | [-0.804, -0.384] | 0.345 |
|  | CETP*Sex | 66 | -0.216 | 0.023 | 0.342 | [-0.666, 0.234] | 0.015 |
| **L**<br>**(TGM2)** | CETP | 66 | -0.315 | 0.014 | 0.035 | [-0.557, -0.073] | 0.100 |
|  | CETP*Sex | 66 | -0.091 | 0.027 | 0.745 | [-0.614, 0.433] | 0.002 |
| **M**<br>**(PPP1R9B)** | CETP | 66 | 0.743 | 0.096 | <0.001 | [0.574, 0.912] | 0.558 |
|  | CETP*Sex | 66 | 0.045 | 0.208 | 0.807 | [-0.321, 0.410] | 0.001 |
| **N**<br>**(WWC1)** | CETP | 66 | -0.493 | 0.470 | <0.001 | [-3.113, -1.234] | 0.260 |
|  | CETP*Sex | 66 | -0.266 | 1.004 | 0.248 | [-0.721, 0.190] | 0.022 |

| **Figure 6** | **Simple Effects** | **Sex** | **N** | **$\beta$** | **SE** | **p** | **$p\eta^2$** |
| --- | --- | --- | --- | --- | --- | --- | --- |
| **A**<br>**(VCIS)** | CETP | Male | 19 | 0.270 | 0.004 | 0.241 | 0.025 |
|  |  | Female | 42 | -0.370 | 0.003 | 0.028 | 0.086 |
| **B**<br>**(VCIS)** | Aβ42 | Male | 15 | 0.372 | 0.000 | 0.195 | 0.044 |
|  |  | Female | 30 | 0.186 | 0.000 | 0.431 | 0.016 |
| **C**<br>**(VCIS)** | p-tau(181)/Aβ42<br>($\times 10^2$) | Male | 15 | -0.495 | 0.015 | 0.047 | 0.099 |
|  |  | Female | 30 | -0.417 | 0.013 | 0.067 | 0.086 |
| **D**<br>**(TAGLN)** | CETP | Male | 20 | -0.434 | 285.377 | 0.014 | 0.097 |
|  |  | Female | 46 | -0.696 | 196.510 | <0.001 | 0.368 |
| **E**<br>**(PECAM1)** | CETP | Male | 20 | 0.541 | 0.025 | <0.001 | 0.182 |
|  |  | Female | 45 | 0.770 | 0.011 | <0.001 | 0.464 |
| **F**<br>**(VEGFR1)** | CETP | Male | 20 | -0.011 | 1.364 | 0.957 | 0.000 |
|  |  | Female | 46 | -0.425 | 0.939 | 0.004 | 0.132 |
| **G**<br>**(VEGFR2)** | CETP | Male | 20 | -0.476 | 22.189 | 0.005 | 0.127 |
|  |  | Female | 46 | -0.785 | 15.280 | <0.001 | 0.454 |
| **H** | CETP | Male | 20 | -0.662 | 0.018 | <0.001 | 0.209 |
| **(CDH5)** |  | Female | 45 | -0.657 | 0.013 | <0.001 | 0.323 |
| **I**<br>**(MMRN2)** | CETP | Male | 20 | -0.215 | 0.086 | 0.313 | 0.017 |
|  |  | Female | 45 | 0.398 | 0.059 | 0.008 | 0.112 |
| **J**<br>**(ATP2A3)** | CETP | Male | 20 | -0.087 | 0.315 | 0.675 | 0.003 |
|  |  | Female | 44 | -0.422 | 0.227 | 0.006 | 0.122 |
| **K**<br>**(CD59)**<br>**(Robust errors)** | CETP | Male | 20 | -0.448 | 0.019 | 0.019 | 0.089 |
|  |  | Female | 46 | -0.663 | 0.013 | <0.001 | 0.311 |
| **L**<br>**(TGM2)** | CETP | Male | 20 | -0.254 | 0.013 | 0.071 | 0.053 |
|  |  | Female | 46 | -0.344 | 0.021 | 0.124 | 0.039 |
| **M**<br>**(PPP1R9B)** | CETP | Male | 20 | 0.713 | 0.171 | <0.001 | 0.272 |
|  |  | Female | 46 | 0.757 | 0.118 | <0.001 | 0.471 |
| **N**<br>**(WWC1)** | CETP | Male | 20 | -0.313 | -1.379 | 0.100 | 0.044 |
|  |  | Female | 46 | -0.579 | -2.550 | <0.001 | 0.251 |

**Supplemental Table 4:** Supplemental Figure 10 detailed statistical analyses.

| **Suppl Figure 10** | **Predictor/Interaction** | **N** | **$\beta$** | **SE** | **p** | **$\beta$ 95% CI** | **$p\eta^2$** |
| --- | --- | --- | --- | --- | --- | --- | --- |
| **A**<br>**(RBANS total)** | p-tau(181)/Aβ42 ($\times 10^2$) | 46 | -0.425 | 0.007 | 0.017 | [-0.771, -0.079] | 0.134 |
| | p-tau(181)/Aβ42 ($\times 10^2$) *Sex | 46 | 0.169 | 0.013 | 0.582 | [-0.447, 0.784] | 0.008 |
| **B**<br>**(RBANS total)** | CETP | 63 | -0.132 | 0.002 | 0.328 | [-0.399, 0.135] | 0.017 |
|  | CETP*Sex | 63 | -0.386 | 0.004 | 0.171 | [-0.012, 0.002] | 0.033 |
| **C**<br>**(RBANS total)** | Aβ42 | 46 | 0.132 | 0.000 | 0.468 | [-0.232, 0.493] | 0.013 |
| | Aβ42*Sex | 46 | -0.088 | 0.000 | 0.814 | [-0.843, 0.666] | 0.001 |
| **D, F**<br>**(VCIS)** | p-tau(181)/Aβ42 ($\times 10^2$) *CETP | 45 | 0.040 | 0.094 | 0.896 | [-0.575, 0.655] | 0.000 |
| | p-tau(181)/Aβ42 ($\times 10^2$) *Sex*CETP | 45 | -0.542 | 0.207 | 0.425 | [-1.905, 0.822] | 0.019 |
| **E, G**<br>**(RBANS total)** | p-tau(181)/Aβ42 ($\times 10^2$) | 46 | 0.091 | 0.013 | 0.760 | [-0.508, 0.691] | 0.002 |
| | p-tau(181)/Aβ42 ($\times 10^2$) *Sex*CETP | 46 | 1.253 | 0.030 | 0.075 | [-0.131, 2.638] | 0.088 |
| **H**<br>**(PPP1R10)**<br>**(Robust SE)** | CETP | 66 | 0.238 | 0.013 | 0.077 | [-0.013, 0.489] | 0.055 |
|  | CETP*Sex | 66 | -0.049 | 0.024 | 0.840 | [-0.592, 0.494] | 0.001 |
| **I**<br>**(TNFSF10)** | CETP | 66 | -0.268 | 0.092 | 0.031 | [-0.512, -0.025] | 0.074 |
|  | CETP*Sex | 66 | 0.666 | 0.187 | 0.010 | [0.169, 1.164] | 0.107 |
| **J**<br>**(OSMR)** | CETP | 66 | -0.227 | 0.116 | 0.074 | [-0.476, 0.023] | 0.053 |
|  | CETP*Sex | 66 | 0.185 | 0.245 | 0.485 | [-0.341, 0.710] | 0.008 |
| **K**<br>**(CETP)** | OligoAlbumin<br>CSF/Plasma Ratio<br>(x10<sup>3</sup>) | 34 | -0.027 | 0.878 | 0.892 | [-0.436, 0.377] | 0.001 |
| **L**<br>**(CETP)** | IgG CSF/Plasma Ratio<br>(x10<sup>3</sup>) | 28 | -0.080 | 1.657 | 0.693 | [-0.494, 0.334] | 0.007 |

| Suppl<br>Figure 10 | Simple Effects | Sex |  | N | β | SE | p | pn<sup>2</sup> |
| --- | --- | --- | --- | --- | --- | --- | --- | --- |
| A | p-tau(181)/Aβ42<br>(x10<sup>2</sup>) | Male |  | 15 | -0.519 | 0.011 | 0.038 | 0.106 |
|  |  | Female |  | 31 | -0.350 | 0.010 | 0.119 | 0.061 |
| B | CETP | Male |  | 15 | 0.127 | 0.003 | 0.580 | 0.005 |
|  |  | Female |  | 31 | -0.259 | 0.002 | 0.114 | 0.044 |
| C | Aβ42 | Male |  | 15 | 0.185 | 0.000 | 0.524 | 0.011 |
|  |  | Female |  | 31 | 0.096 | 0.002 | 0.687 | 0.004 |
| D, F | p-tau(181)/Aβ42<br>(x10<sup>2</sup>) | Male | <50%<br>CETP | 7 | -0.429 | 0.085 | 0.130 | 0.066 |
|  |  |  | >50%<br>CETP | 8 | -0.365 | 0.141 | 0.428 | 0.019 |
|  |  | Female | <50%<br>CETP | 17 | -0.009 | 0.118 | 0.981 | 0.000 |
|  |  |  | >50%<br>CETP | 13 | -0.487 | 0.076 | 0.055 | 0.104 |
| E, G | p-tau(181)/Aβ42<br>(x10<sup>2</sup>) | Male | <50%<br>CETP | 7 | -0.381 | 0.012 | 0.185 | 0.050 |
|  |  |  | >50%<br>CETP | 8 | -1.160 | 0.020 | 0.017 | 0.151 |
|  |  | Female | <50%<br>CETP | 17 | -0.690 | 0.017 | 0.085 | 0.082 |
|  |  |  | >50%<br>CETP | 14 | -0.216 | 0.011 | 0.388 | 0.021 |
| H | CETP | Male |  | 20 | 0.271 | 0.018 | 0.128 | 0.038 |
|  |  | Female |  | 46 | 0.222 | 0.018 | 0.212 | 0.026 |
| I | CETP | Male |  | 20 | -0.720 | 0.154 | <0.001 | 0.171 |
|  |  | Female |  | 46 | -0.054 | 0.106 | 0.704 | 0.002 |
| J | CETP | Male |  | 20 | -0.346 | 0.197 | 0.107 | 0.044 |
|  |  | Female |  | 44 | -0.162 | 0.145 | 0.302 | 0.018 |

